# LNA043 and ANGPTL3 Interactions with Integrin α5β1 and Fibronectin Drive Distinct Stromal and Immune Responses

**DOI:** 10.64898/2026.09.14.751366

**Authors:** Andrea Grosso, Stefano Cucuzza, Elsa Gambs, Thomas Lapointe, Truc Huynh, Guray Kuzu, Frederik King, Kenneth Ng, Jian Shi, Markus Vogel, Nicole Gerwin, Christine Halleux, Mara Fornaro

## Abstract

Osteoarthritis (OA) is characterized by cartilage breakdown, extracellular matrix remodeling, and synovial inflammation. Dysregulated integrin α5β1 signaling and fibronectin fragmentation are key drivers of OA progression.

We found that in ihMSC, LNA043, Full length ANGPTL3 (FL-ANGPTL3), and C-terminal fibrinogen like domain of ANGPTL3 (C-ANGPTL3) interacted with both integrin α5β1 and fibronectin, consistent with a multivalent binding mode supported by AlphaFold based structural predictions. FL-ANGPTL3 supported ihMSC adhesion and induced human monocyte migration, with both effects being dependent on integrin β1. C-ANGPTL3 alone was sufficient to mediate these effects, although with reduced efficacy compared to the full-length protein. In contrast, LNA043 did not support ihMSC adhesion but retained integrin β1-dependent monocyte migratory activity, defining a more selective functional profile and indicating that further truncation modulates downstream responses without abolishing integrin engagement. To further explore the biology of ANGPTL3 and the mechanism of action of LNA043 we employed a targeted RASL Seq profiling. We found that both FL- and C-ANGPTL3, but not LNA043, induced broad transcriptional reprogramming and counter regulated fibronectin fragment–driven inflammatory signaling in monocytes.

We uncovered a role for ANGPTL3 in fibronectin-integrin α5β1 signaling and showed that LNA043 preserves this interaction while uncoupling it from strong immune responses.

## Introduction

Osteoarthritis (OA) is the most common degenerative joint disease worldwide and a leading cause of disability among the elderly population (1). It is characterized by progressive degradation of cartilage tissue, chronic low-grade inflammation, and extensive remodeling of the extracellular matrix (ECM) (2,3). While the loss of articular cartilage is a hallmark of OA, it is now established that the synovial microenvironment, including resident and infiltrating immune cells, plays a central role in driving disease progression and joint dysfunction (4). Among these, monocytes and macrophages are key mediators of inflammation and tissue repair, orchestrating responses to matrix-derived damage-associated molecular patterns (DAMPs) that accumulate in diseased joints (5–8). A prominent ECM alteration during OA progression is the proteolytic breakdown of fibronectin, which generates biologically active fragments that profoundly influence cellular behavior (9,10). These fibronectin fragments act as potent DAMPs and activate integrin-dependent signaling pathways not only in chondrocytes, mesenchymal stem cells (MSCs) and synovial cells, but also in monocytes and macrophages (11–13), thereby linking matrix degradation to inflammatory activation of both stromal and immune cells.

Integrin α5β1 is a key component of this complex network. Binding of Fibronectin type III repeats 8–10 (FNIII 8–10) to integrin α5β1 induce the expression of chemokines, cytokines and matrix-remodeling enzymes, as well as cytoskeletal remodeling and metabolic reprogramming (14,15). Immune cells such as monocytes and macrophages are highly sensitive to these signals, adopting inflammatory phenotypes that exacerbate tissue damage and impair regeneration. Modulating α5β1 integrin-driven responses in chondrocytes, stromal and immune compartments represent a potential therapeutic approach for OA. In this context, LNA043, an engineered fragment of the fibrinogen-like domain of angiopoietin-like 3 (ANGPTL3), has emerged as a promising disease-modifying OA treatment candidate targeting integrin α5β1(16). However, despite its regenerative potential, the molecular mechanism of LNA043 remains elusive.

The angiopoietin-like (ANGPTL) protein family consists of secreted proteins that play roles in angiogenesis, metabolic processes, and tissue remodeling. Different ANGPTL members, including ANGPTL2, ANGPTL3 and ANGPTL4, are known to regulate immune cell trafficking and macrophage activation (17–19), suggesting an important role of this protein family in ECM biology and inflammation. ANGPTL3, primarily produced by the liver, has long been recognized as a hepatokine regulating lipid metabolism, cardiovascular risk, and HDL function (20–23). Structurally, ANGPTL3 contains an N-terminal coiled-coil domain responsible for inhibiting lipoprotein lipase (24), and a C-terminal fibrinogen-like domain (FBN) that mediates additional biological activities. Through its FBN domain, ANGPTL3 promotes endothelial adhesion, proliferation, and migration via αVβ3 integrin and participates in vascular inflammation (25). Recent studies showed that ANGPTL3 modulates monocyte and macrophage behavior, regulating the expression of cytokines such as IL-1β, IL-6, and TNFα through integrin-dependent mechanisms (26,27).

Here, we investigated how LNA043, full-length ANGPTL3 (FL-ANGPTL3), and the C-terminal FBN domain of ANGPTL3 (C-ANGPTL3) engage integrins and if these interactions regulate MSC adhesion as well as monocyte migration and transcriptional responses. We then assessed their ability to counter-act FNIII 8–10–induced inflammation in monocytes. Collectively, our study aims to elucidate how LNA043 and ANGPTL3 modulate key integrin biological activity, and to define how these interactions contribute to their immunomodulatory and regenerative potential in OA.

## Results

### LNA043 Co-localizes with Integrin α5β1 and Fibronectin at Fibrillar Adhesions in ihMSC

ANGPTL3 has been shown to interact with the integrin αVβ3 in endothelial cells (25). However, in a previous study, we found that LNA043 binds specifically to the integrin α5β1 (16). To confirm and better understand the interaction of LNA043 with integrin α5β1, we treated the immortalized human bone marrow-derived mesenchymal stem cell line UE7T-13 (ihMSC) with LNA043 for 1 hour. Immunofluorescence staining of ihMSCs treated with LNA043 revealed strong co-localization of LNA043 with both α5β1 integrin and fibronectin (Figure 1A). Higher-magnification orthogonal confocal views (Figure 1B) showed that LNA043 was present along fibrillar adhesions, structures that connect cells to fibronectin fibrils through integrin α5β1 (28). Similar to LNA043, both the native FL-ANGPTL3 and C-ANGPTL3 strongly colocalized with integrin α5β1 and fibronectin (Figure S1A). Notably, we found that LNA043, as well as FL-ANGPTL3 and C-ANGPTL3, did not significantly colocalize with integrin αVβ3 (Figure S1B), which was primarily found at focal adhesion sites.

**Figure 1.**
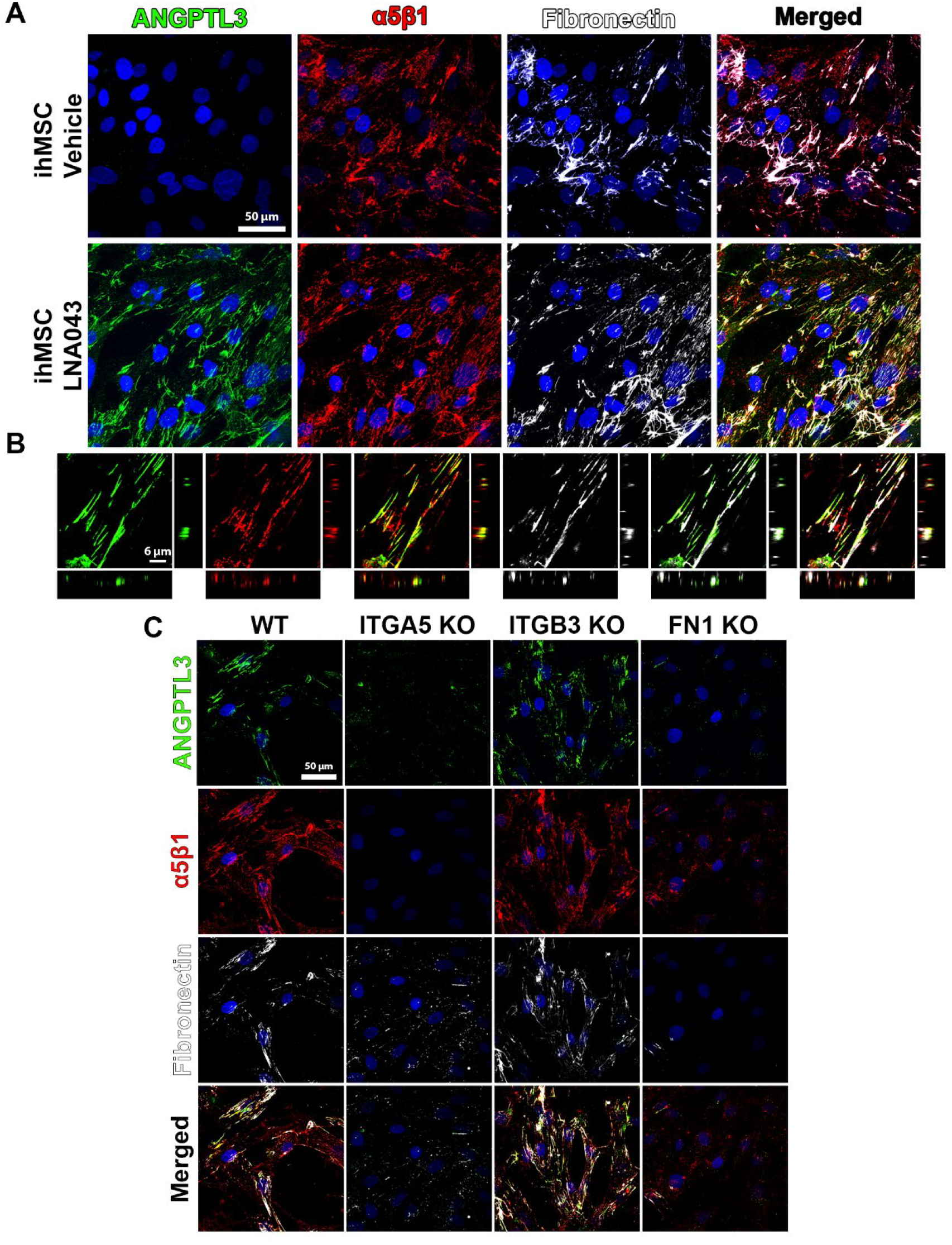
Localization of LNA043 in ihMSCs following in vitro treatment. (A) Immunofluorescent staining of ihMSCs after 1 hour treatment with vehicle control (top panels) or 300 µg/mL (12 µM) LNA043 (bottom panels), showing LNA043 (stained with an anti-ANGPTL3 antibody, green), integrin α5β1 (red), and cellular fibronectin (white). (B) High magnification orthogonal view of fibrillar adhesion structures illustrating co-localization of LNA043 with both integrin α5β1 and fibronectin. (C) Immunofluorescent staining of wild-type, ITGA5 knockout, ITGB3 knockout, and FN1 knockout ihMSCs after 1 hour treatment with 300 µg/mL (12µM) LNA043. Scale bars: A = 50 µm; B = 6 µm; C = 50 µm.

To assess whether integrin α5β1 or fibronectin are required for LNA043 interaction with ihMSCs, we knocked out integrin α5, β3, and fibronectin (FN1) in ihMSC cells and assessed LNA043 localization by immunofluorescence after 1 hour of treatment. LNA043 was nearly undetectable at the cell surface in both α5 and FN1 knockout cells, in which the loss of α5 or fibronectin prevents fibrillar adhesion assembly and thereby LNA043 binding. LNA043 localization, instead, remained unchanged in integrin β3 knockout cells compared to wild type cells (Figure 1C). These results suggest that both fibronectin and α5β1, but not αVβ3 integrin, are essential for LNA043 binding to the cell surface and its incorporation into the extracellular matrix.

### LNA043, C- and FL-ANGPTL3 Selectively Bind the Heparin-Binding Domain II of Fibronectin

The integrin α5β1–fibronectin axis is frequently dysregulated in OA, contributing to matrix degradation and chondrocyte hypertrophy (14,29–31). We previously showed that LNA043 preferentially penetrates extensively degraded fibronectin-rich cartilage, suggesting that LNA043 can be incorporated into the cartilage ECM (16). Fibronectin is composed of multiple functional domains (Figure 2A), among which the region encompassing its 12th to 14th type III repeats (FNIII 12–14) is the heparin binding domain II (HBDII), a highly promiscuous growth factor-binding domain that regulates growth factor availability and spatial distribution within tissues (32,33). Instead, integrin α5β1 binds to the RGD motif in FNIII-10 and the synergy site in FNIII-9 and is involved in mediating cell–matrix interactions and chondrogenic signaling (34).

**Figure 2.**
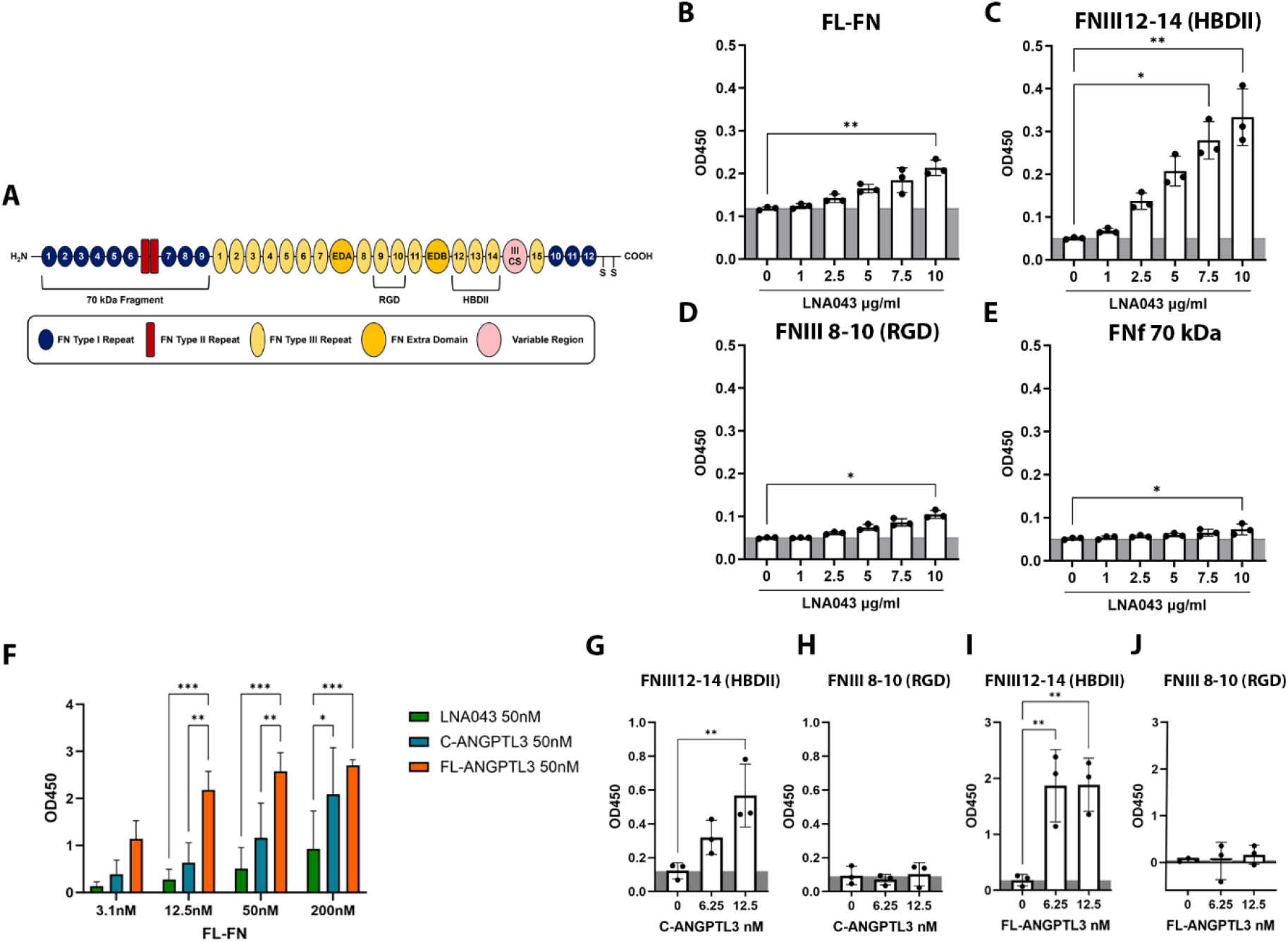
ELISA-based analysis of binding interactions of LNA043, C- and FL-ANGPTL3 with human fibronectin. (A) Schematic illustrating the structure of human FN. (B-E) Concentration-dependent binding of LNA043 to 50 nM of immobilized FL-FN (B), FNIII 12-14 (C), FNIII 8-10 (D) and the 70kDa fragment (E). (F) Comparative analysis of the binding of increasing concentrations of FN-FL to 50 nM LNA43, C-or FL-ANGPTL3. (G-J) Binding profiles of C-ANGPTL3 and FL-ANGPTL3 to 50 nM FNIII 12-14 and FNIII 8-10. Data are shown as mean ± SD from 3 independent experiments. *p < 0.05. **p < 0.01, ***p < 0.001.

We initially assessed binding of LNA043 to human fibronectin using an indirect ELISA. We coated 96-well plates with increasing concentrations of human plasma fibronectin (FL-FN), followed by 1 hour incubation with LNA043 and confirmed that LNA043 binds to FL-FN in a concentration-dependent manner (Figure 2B). To identify the specific FN domain involved in this interaction, we generated the FNIII 12-14 and the FNIII 8-10 fragments and obtained the distal 70kDa N-terminal domain from a commercial source. We found that LNA043 specifically binds to FNIII 12–14, with negligible interaction with the other FN fragments (Figure 2C-E). C-ANGPTL3, and FL-ANGPTL3, could bind FL-FN, with FL-ANGPTL3 showing the highest binding (Figure 2F). Similarly to LNA043, both C-ANGPTL3 and FL-ANGPTL3 primarily interacted with the HBDII domain (Figure 2G-J), supporting a role of this region in mediating fibronectin interaction with ANGPTL3 and its derived truncated proteins.

### LNA043 Simultaneously Engages Integrin α5β1 and Fibronectin: Structural Predictions and Experimental Validation of a Multi-Partner Binding Model

Co-localization and binding experiments suggested that LNA043, C- and FL-ANGPTL3, interact with both integrin α5β1 and FN. To gain deeper structural insight into those interactions, we generated AlphaFold models of the respective complexes and performed molecular dynamics (MD) simulations. We then used short- and long-range interaction energies as proxies to estimate relative binding affinities.

Structural predictions showed that LNA043 interacts with integrin α5β1 headpieces through its predominantly unstructured C-terminal region (Figure 3A). Modeling of multiple αβ integrin heterodimers yielded similar complex relative orientations, with the unstructured C-terminal region of LNA043 engaging the interface between α and β headpieces (Figure S2B). Interestingly, the predicted binding interface of LNA043 with integrin α5β1 appears to resemble the RGD-binding motif present in FN, both in orientation and structure (Figure 3A, inset). This suggests that LNA043 might engage integrin α5β1 through a mechanism that partially mimics the RGD-binding mode of FN. However, the different primary sequence of the LNA043 loop likely results in differences in binding affinity and/or specificity.

**Figure 3.**
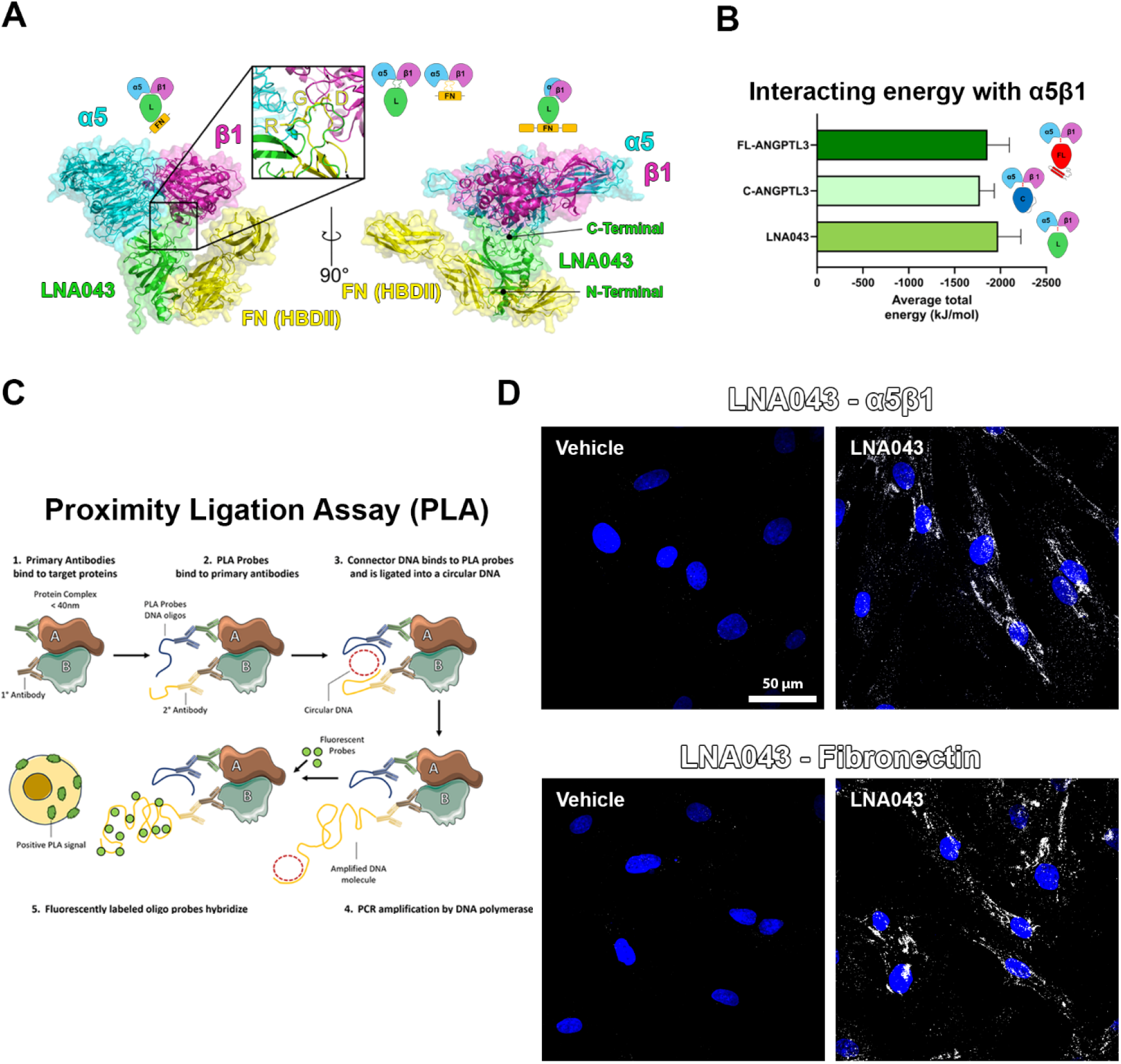
Structural Predictions and Experimental Validation of Multi-Partner Binding Model. (A) AlphaFold and molecular dynamics simulations showing the predicted dual binding of LNA043 to integrin α5β1 via its unstructured C-terminal region and to the HBDII domain of fibronectin via its N-terminal region. Integrin α5 subunit is shown in cyan, β1 subunit in magenta, LNA043 in green, and HBDII in yellow. The inset highlights the overlap between the unstructured loop of LNA043 and the RGD loop of fibronectin. (B) Total average interacting energy between α5β1 integrin and LNA043, C-ANGPTL3 and FL-ANGPTL3. The root mean square deviation (RMSD) of the total energy is shown as error bars. (C) Schematic representation of proximity ligation assay (PLA) used to detect close spatial proximity between LNA043 and integrin α5β1 or fibronectin in ihMSCs. (D) Representative PLA images of ihMSCs treated with vehicle (left panel) or 300 μg/ml LNA043 (right panel), confirming proximity of LNA043 to integrin α5β1 (top right panel) or fibronectin (bottom right panel). Positive PLA signal is shown in white. Scale bar = 50 µm.

To further investigate potential interactions, we modeled the complex between LNA043 and the FN HBDII fragment. Three binding conformations were predicted, involving the N-Terminal, central, or C-terminal regions of LNA043 (Figure S2C). Among these, the conformation involving the N-terminal region (conformation “3”) showed the lowest average total interaction energy, suggesting it is the most likely to occur. Binding via the C-terminal region of LNA043 would compete with the predicted integrin-binding interface, which showed a more favorable interaction energy (Supplementary Table). These data support the hypothesis that LNA043 is able to interact with both integrin α5β1 and fibronectin using spatially distinct regions. This dual engagement could enable the formation of a ternary FN:LNA043:α5β1 complex, in which the interaction between LNA043 and integrin α5β1 is predicted to be the most energetically favorable (Supplementary Video). In line with these findings, proximity ligation assay (PLA) performed in ihMSCs treated with LNA043 revealed close proximity of LNA043 to both fibronectin and integrin α5β1, consistent with the formation of a ternary FN:LNA043:α5β1 complex in a cellular context (Figure 3C–D). In silico modeling of C-ANGPTL3 and FL-ANGPTL3 showed similar orientations (Figure S2D) and interaction energies (Figure 3B), indicating that the C-terminal region of ANGPTL3 is sufficient for binding to both fibronectin and integrin α5β1. Additional modeling of C- and FL-ANGPTL3 with the FN HBDII revealed that while the C-terminal interaction was conserved, the N-terminal binding solutions were absent (Figure S2E). This may indicate that the additional amino acids at the N-terminus hinder that binding interface. Importantly, this data do not rule out an interaction between C- or FL-ANGPTL3 and fibronectin but instead suggest a shift in binding equilibrium toward the C-terminal region.

### FL- and C-ANGPTL3 promote ihMSC adhesion in a dose- and β1 integrin-dependent manner, unlike LNA043

Integrins are key mediators of cell adhesion, anchoring cells to the extracellular matrix and transmitting signals that regulate survival, migration, and differentiation (35,36). ANGPTL3 has been reported to promote endothelial cell adhesion through integrin αVβ3 (25). To investigate whether LNA043 was able to support cell adhesion, we performed adhesion assays using ihMSC. Plates were coated with increasing concentrations of LNA043 (0-2 µM) and ihMSC were seeded onto the coated plates for 1 hour in serum free medium containing 1mM of CaCl_2_, MgCl_2_ and 0.5 mM of MnCl_2_. As shown in Figure 4A, LNA043 did not promote cell adhesion at any of the concentrations tested. We next assessed whether ANGPTL3 supported ihMSCs adhesion as previously reported in endothelial cells. Immortalized hMSC were seeded on wells coated with FL-ANGPTL3 or C-ANGPTL3 at the same concentration range (0-2 μM) used for LNA043. Both proteins induced a robust, concentration-dependent increase in adhesion, with FL-ANGPTL3 exhibiting a greater effect compared to C-ANGPTL3 (Figure 4A, C).

**Figure 4.**
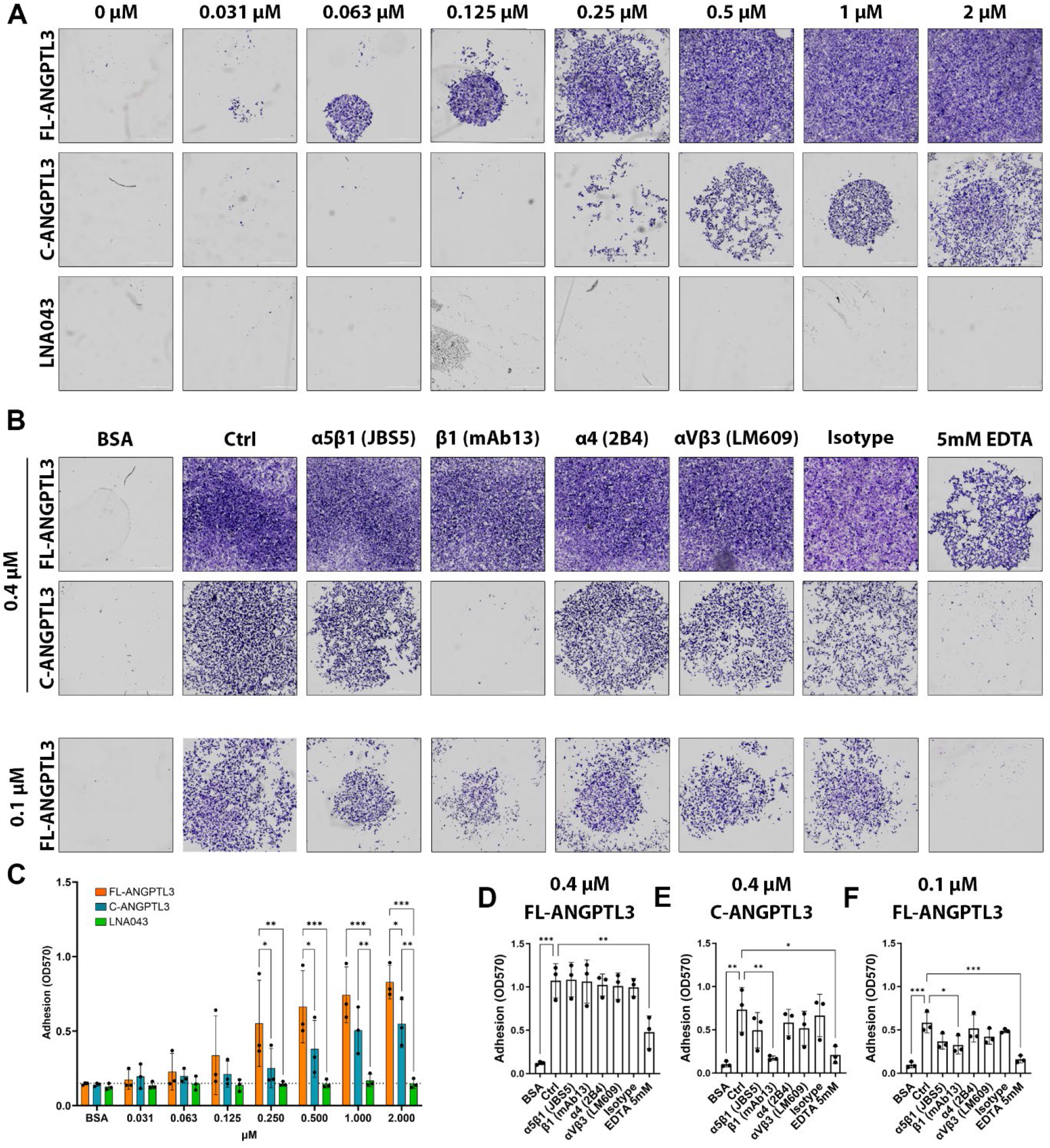
Adhesion of ihMSCs to LNA043, C- and FL-ANGPTL3 and integrin dependency. (A) Representative images of ihMSCs adherent to wells coated with increasing concentrations of LNA043, FL-ANGPTL3, or C-ANGPTL3 (0–2 µM) after 1 hour at 37 °C. (B) Representative images of ihMSCs adherent to 0.4 µM FL-ANGPTL3 or C-ANGPTL3 (top) and 0.1 µM FL-ANGPTL3 (bottom) following pre-incubation with integrin-blocking antibodies (α5β1, β1, α4, αVβ3), isotype controls, or 5 mM EDTA. (C) Quantification of dose-dependent adhesion (panel A). (D–F) Quantification of adhesion after integrin inhibition (panel B). Data are shown as mean ± SD from 3 independent experiments. *p < 0.05. **p < 0.01, ***p < 0.001.

To elucidate the molecular mechanisms underlying ANGPTL3-mediated adhesion, we examined the involvement of integrins using function blocking antibodies and EDTA, which chelates divalent cations (Ca²⁺ and Mg²⁺) essential for integrin activation and ligand binding. Cells were pre-treated for 30 minutes with blocking antibodies targeting α5β1 (clone JB55), β1 (clone mAb13), α4 (clone 2B4), or αVβ3 (clone LM609), isotype controls or 5 mM EDTA. Cells were then seeded for 1 hour on plates coated with 0.4 μM FL-ANGPTL3, 0.4uM C-ANGPTL3, or 0.1 μM FL-ANGPTL3 (Figures 4B, 4D–F). Pre-treatment with mAb13, a β1 integrin-blocking antibody, completely abolished ihMSC adhesion to 0.4 µM C-ANGPTL3. In contrast, none of the antibodies affected adhesion to 0.4 µM FL-ANGPTL3. When the coating concentration of FL-ANGPTL3 was reduced to 0.1 µM, pre-treatment with the β1-blocking antibody significantly decreased, but did not fully prevent cell adhesion.

Since LNA043 was produced in CHO cells and differs from C-ANGPTL3 by a single point mutation (K423Q) and a 17-amino acid (aa) deletion at the N-terminus, we investigated whether the production system, potential differences in glycosylation, the 17-aa deletion, or the point mutation could influence its ability to promote adhesion. To this end, we used the same expression system used for FL- and C-ANGPTL3, and generated LNA043 in Expi293 cells, with and without the K423Q mutation. Adhesion assays revealed that neither the production system nor the point mutation restored the adhesion-promoting activity, indicating that these factors do not account for the lack of cell adhesion observed with LNA043 (Figure S3A, B). These findings suggest that the 17-aa stretch missing in LNA043 likely plays an important role in mediating adhesion. Collectively, these results demonstrate that ANGPTL3 supports cell adhesion in an integrin-dependent manner and that the C-terminal FBN domain alone is sufficient to mediate this effect.

### LNA043 induces THP-1 cell migration in a dose and integrin β1-dependent manner

In our previous studies we showed that LNA043 induces chondrogenesis, preserves cartilage matrix protein expression, and exerts anti-inflammatory effects on human MSCs and chondrocytes in vitro (16). Given that OA is not only a cartilage disease but also affects the synovial environment, these findings raised the question of whether LNA043 could also modulate inflammatory cells within other joint tissues. The synovial membrane often exhibits low-grade chronic inflammation, where resident macrophages and monocytes, whether infiltrating or locally present, influence both inflammatory responses and tissue remodeling (37–39). Understanding how LNA043 interacts with immune cells is critical for evaluating its potential immunomodulatory and cartilage repair effects. Monocyte migration is a critical step in inflammation and tissue repair. Circulating monocytes are recruited to sites of injury or degeneration, where they differentiate into macrophages and contribute to matrix remodeling, angiogenesis, and immune regulation (40). Notably, other members of the ANGPTL family, such as ANGPTL2 and ANGPTL4, have been implicated in immune cell trafficking, and modulation of inflammatory processes (17,18,31,41,42) supporting the hypothesis that LNA043 might exert similar effects. To determine whether LNA043 influences monocyte migration, we performed transwell migration assays using THP-1, a human monocytic cell line. Cells were seeded in the upper chamber of the transwell and increasing concentrations of LNA043 were added to the lower chamber (Figure 5A). LNA043 induced a dose-dependent increase in THP-1 migration at 6 and 24 hours (Figure 5B). To explore the role of integrins, THP-1 cells were pre-treated for 30 minutes with blocking antibodies targeting α5β1 (clone JB55), β1 (clone mAb13), α4 (clone 2B4), or αVβ3 (clone LM609), as well as isotype controls, prior to performing migration assays with LNA043 (Figure 5C). Blockade of α5β1, α4β1 or β1 significantly reduced migration compared to the isotype control, whereas αVβ3 inhibition had no significant effect (Figure 5D). These findings indicate that LNA043 promotes THP-1 migration primarily through β1-containing integrins. Importantly, similar results were obtained when using LNA043 produced in Expi293 cells, and no significant differences were observed between the wild-type and mutant forms of the protein, indicating that neither the production system nor the introduced mutation impacted the migration-promoting activity of LNA043 (Figure S3C-D).

**Figure 5.**
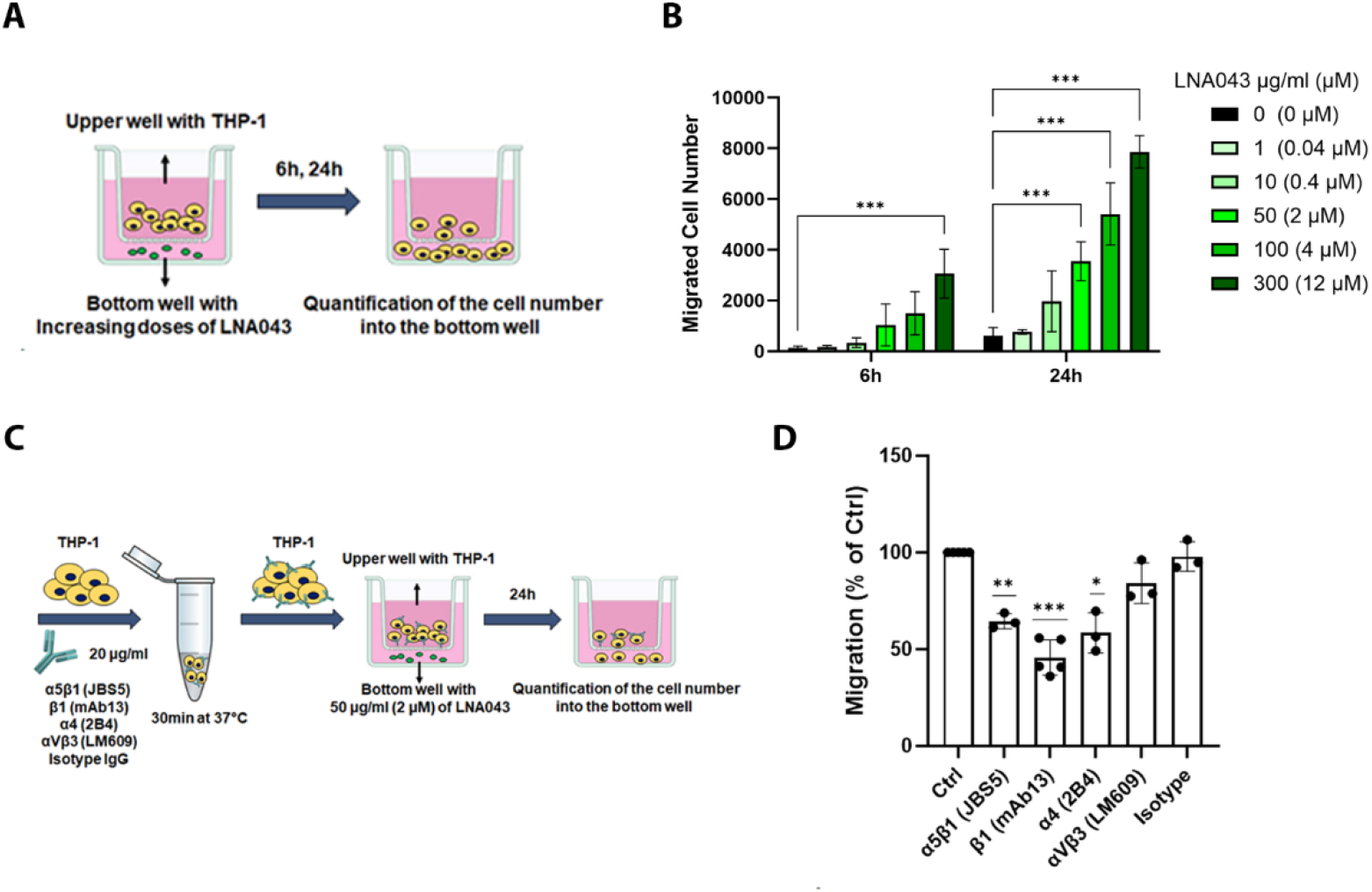
THP-1 transwell migration assay in response to LNA043. (A) Schematic of transwell migration assay. THP-1 cells were seeded in the upper chamber and increasing concentrations of LNA043 were added to the lower chamber. Migration was assessed after 6 hours and 24 hours. (B) Dose-dependent migration of THP-1 cells in response to LNA043 at 6 hours and 24 hours. (C) Experimental design for integrin-blocking assays. THP-1 cells were pre-incubated with antibodies targeting α5β1 (JB55), β1 (mAb13), α4 (2B4), or αVβ3 (LM609), or isotype controls, prior to migration assessment with 2 µM LNA043. (D) Quantification of migration after integrin blockade, expressed as percentage of control. Data are shown as mean ± SD from 3 independent experiments. *p < 0.05. **p < 0.01, ***p < 0.001.

### LNA043 and ANGPTL3 Promote THP-1 and Human Primary Monocyte Migration

Next, we compared the ability of LNA043, C- and FL-ANGPTL3 to promote monocyte migration. After 24 hours treatment, both FL-ANGPTL3 and C-ANGPTL3 significantly increased THP-1 migration compared to LNA043, with FL-ANGPTL3 showing the strongest effect (Figure 6A). To assess whether this effect was integrin-dependent, THP-1 cells were pre-treated with a β1-blocking antibody for 30 minutes. Migration was reduced upon β1 integrin inhibition (Figure 6B), confirming the role of β1-containing integrins. Flow cytometry analysis showed that THP-1 cells express α5β1 but not αVβ3 integrins (Figure 6C), consistent with the lack of effect observed when αVβ3 was blocked. Beyond promoting migration, both FL-ANGPTL3 and C-ANGPTL3, but not LNA043, led to an increase in THP-1 cell numbers after 24 hours. This suggests that these proteins not only attract monocytes but also facilitate their initial proliferation (Figure S4). We then validated these findings in primary human monocytes isolated from three independent donors. Both FL-ANGPTL3 and C-ANGPTL3 significantly enhanced migration at 6 and 24 hours across all donors, with FL-ANGPTL3 consistently inducing the highest response (Figures 6D-E). LNA043 also promoted migration of primary monocytes, but only at higher concentrations and to a lesser extent compared to FL- and C-ANGPTL3 (Figure 6D-E). Flow cytometry confirmed that α5β1 integrin is highly expressed in primary monocytes, whereas αVβ3 expression is very low and variable among donors (Figures 6F, 6G).

**Figure 6.**
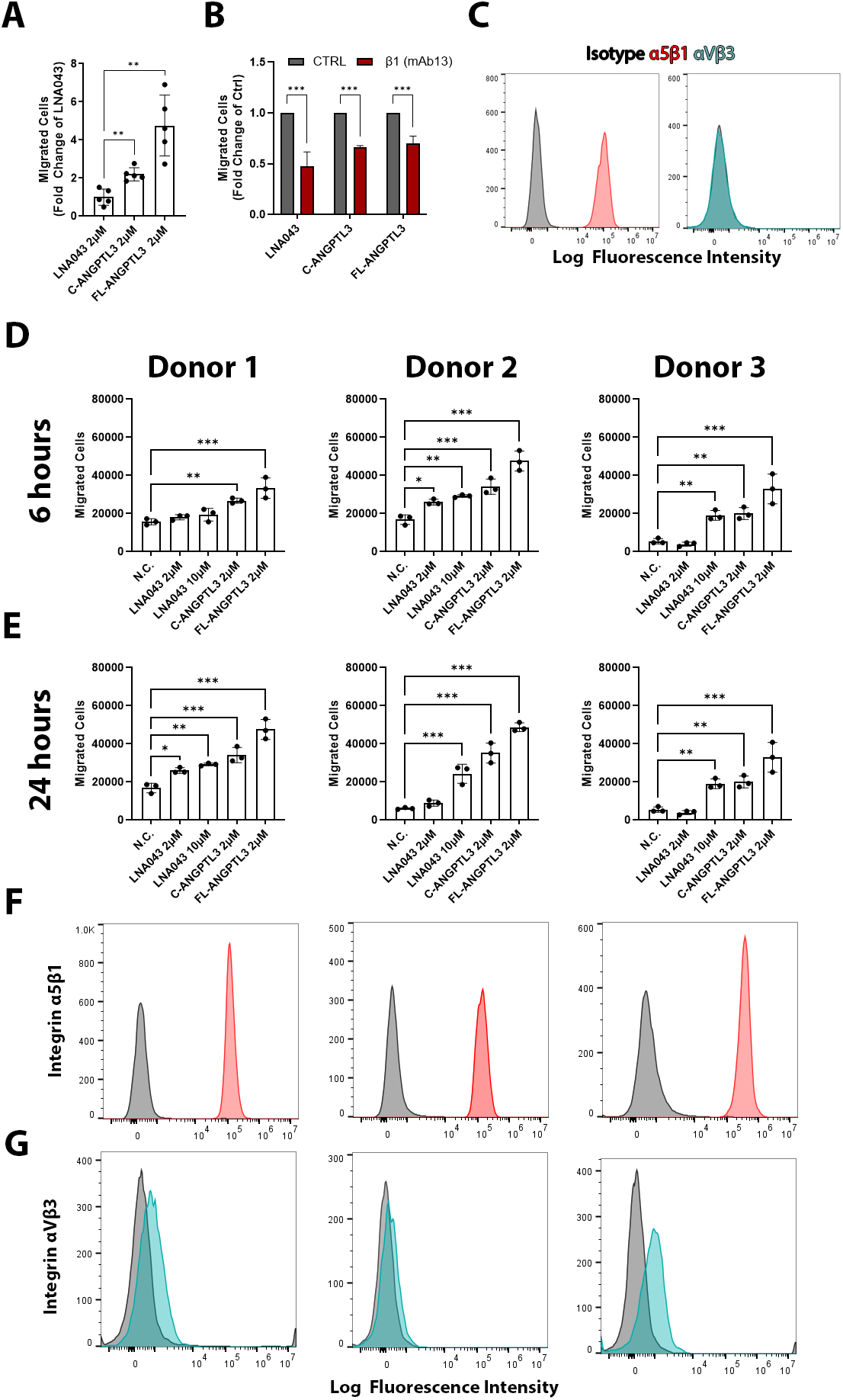
Comparison of migration potency of LNA043, C-ANGPTL3, FL-ANGPTl3 in THP-1 and primary human monocytes. (A) Quantification of THP-1 cell migration toward 2 µM LNA043, C-ANGPTL3, or FL-ANGPTL3 after 24 hours in transwell assays, expressed as fold change of LNA043. (B) Effect of β1 integrin blockade on THP-1 migration toward 2 µM ANGPTL3-derived proteins after 24 hours, expressed as fold change relative to respective controls. (C) Flow cytometry analysis of integrin expression in THP-1 cells (α5β1 in red, αVβ3 in cyan). (D–E) Migration of primary human monocytes from three independent donors toward LNA043, C-ANGPTL3, and FL-ANGPTL3 after 6 (D) and 24 hours (E). (F–G) Flow cytometry analysis of α5β1 and αVβ3 integrin expression in primary monocytes. Data are shown as mean ± SD from 3 independent experiments. *p < 0.05. **p < 0.01, ***p < 0.001.

### FL- and C-ANGPTL3 Drive Robust Transcriptomic Reprogramming in Monocytes, While LNA043 Shows Minimal Effect

To elucidate the mechanisms underlying the effects of LNA043 and ANGPTL3 on THP-1 we performed targeted RASL-Seq profiling following 24 hours treatment with increasing concentrations of LNA043, C-ANGPTL3 or FL-ANGPTL3. The custom RASL-Seq probe panel was designed to analyze 1,764 genes capturing key aspects of immune-regulatory transcriptional programs, including a wide repertoire of chemokines, cytokines, differentiation markers, and genes involved in general cell biology. Both C-ANGPTL3 and FL-ANGPTL3 induced a similar robust, dose-dependent transcriptional regulation, as evidenced by a high number of differentially expressed genes (DEGs) (Figure S5A) and a strong correlation between their gene expression profiles (R²=0.671; Figure 7C). These included upregulation of chemokines (CCL20, CXCL1), extracellular matrix components (FN1), and ECM remodeling enzymes (MMP2), alongside immune regulators (IRF7, HES1, PDPN, AXL, SKIL, BCL2A, NUPR, SPP1), while FABP4, a lipid metabolism marker, was downregulated (Figure 7A-B). Importantly, FABP4 was the only gene that showed significant downregulation by both C- and FL-ANGPTL3 and was also the only gene differentially expressed in response to the highest concentration of LNA043 (Figure S5A). The transcription profile of LNA043-treated THP1 was markedly different from that of C-ANGPTL3, as evidenced by a lower correlation (R²=0.131) (Figure 7D). To validate RASL-Seq findings, we performed qPCR analysis on selected genes after 24 hours treatment with increasing concentrations of LNA043, C-ANGPTL3, or FL-ANGPTL3. Consistent with the transcriptomic data, C-ANGPTL3 and FL-ANGPTL3 induced dose-dependent upregulation of PDPN, SKIL, CCL20, AXL, FN1, MMP2, CXCL1, while downregulating FABP4. Instead, LNA043 only induced a significant, dose-dependent downregulation of FABP4, while all other analyzed genes showed non-significant changes (≤1.5-fold) (Figure 7F). To corroborate these transcriptional changes at the protein level, we quantified by ELISA CCL20, Fibronectin, and MMP2 in culture supernatants and confirmed that C-ANGPTL3 and FL-ANGPTL3 promoted a dose-dependent increase in protein secretion, whereas LNA043 had no measurable effect (Figure 7E).

**Figure 7.**
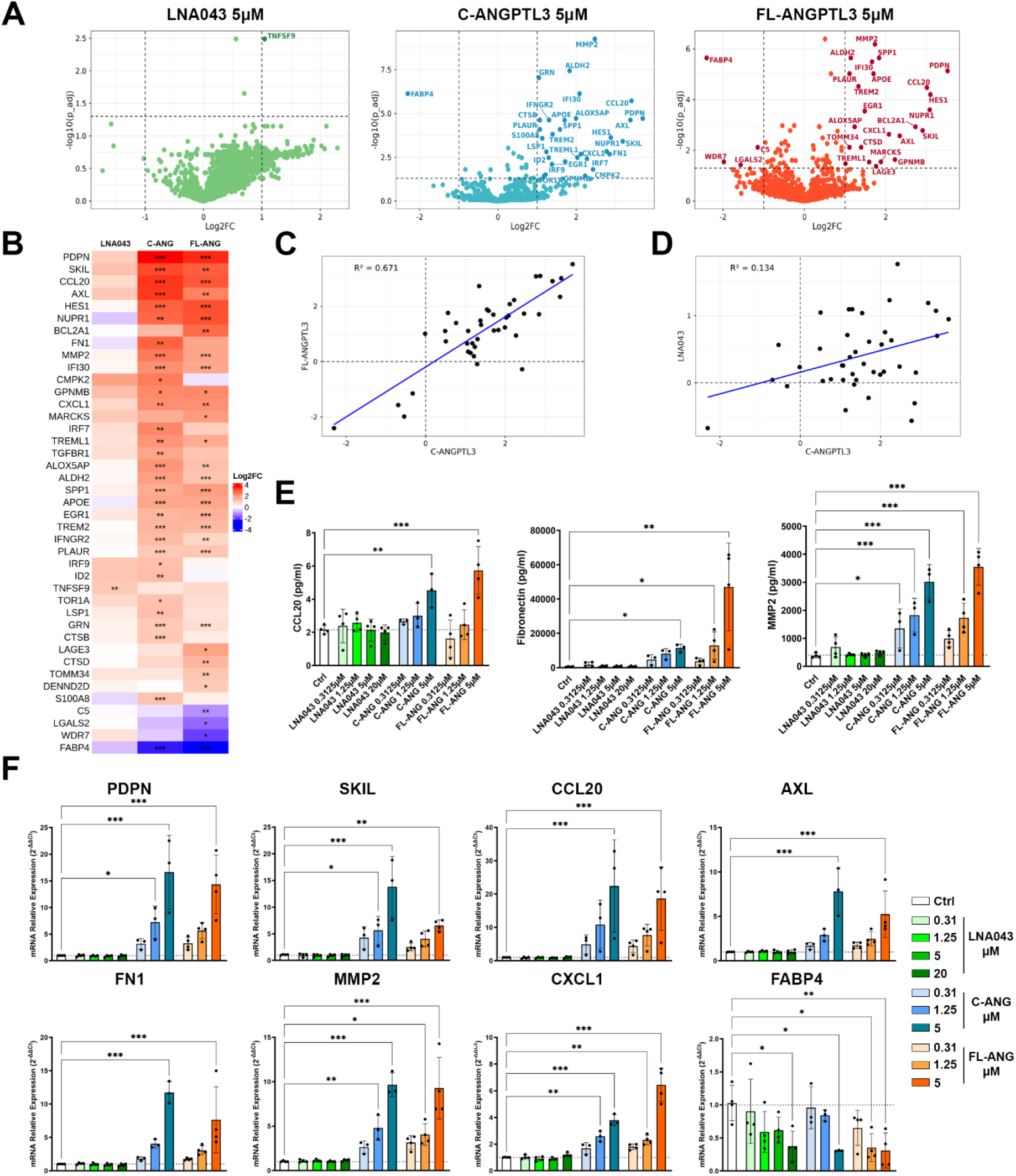
Targeted RASL-Seq transcriptomic analysis of THP-1 treated with LNA043, C-ANGPTL3 and FL-ANGPTL3. (A) Volcano plots illustrating differentially expressed genes (DEGs) after 24 hours treatment with 5 µM of LNA043 (green), C-ANGPTL3 (cyan) or FL-ANGPTL3 (Orange). Data are expressed as log2 fold change (LOG2FC, x-axis) relative to untreated control, and significance is expressed as -log10 adjusted p-value (- log10(p_adj), y axis). Genes are considered significantly regulated above thresholds of LOG2FC ≥ 1 or ≤ –1 and –log10(p_adj) ≥ 1.3 (p_adj < 0.05). (B) Heatmap of significantly regulated genes after 24 hours treatment with 5 µM LNA043, C-ANGPTL3, or FL-ANGPTL3, based on hierarchical clustering of log2 fold change values. Color scale represents relative expression changes, with red indicating upregulation and blue indicating downregulation, while asterisks indicate level of statistical significance. (C,D) Correlation analysis of gene expression profiles on significantly regulated genes between C-ANGPTL3 and FL-ANGPTL3 (C) and between C-ANGPTL3 and LNA043 (D). Each point represents a gene meeting the significance criteria, and the blue linear regression lines indicate the degree of correlation between treatments using Pearson’s correlation. (E) Measurement of CCL20, fibronectin, and MMP2 protein levels in THP-1 cell supernatants by ELISA following 24 hours treatment with increasing concentrations of LNA043, C-ANGPTL3, or FL-ANGPTL3. (F) qPCR validation of top representative genes after 24 hours treatment with increasing concentrations of LNA043, C-ANGPTL3, or FL-ANGPTL3. Data are shown as mean ± SD from 3-4 independent experiments. *p < 0.05, **p < 0.01, ***p < 0.001.

### FL-ANGPTL3 Counteracts FNIII 8-10 Induced Inflammatory Activation and Promotes a Pro-Resolutive Transcriptomic Shift in THP-1 monocytes

To determine whether the immunoregulatory phenotype induced by ANGPTL3 could shift monocytes toward a pro-resolutive state in the context of joint inflammation, we sought to model the inflammatory microenvironment characteristic of OA. Since the FNIII 8-10 fragment is abundantly present in OA joints and functions as a matrix-derived DAMP via integrin α5β1 (9,15), we used it to mimic the inflammatory cues encountered by monocytes in diseased tissue. RASL-Seq profiling of THP-1 treated with 1 μM FNIII 8-10 for 24 hours showed a pronounced pro-inflammatory transcriptional program activation (Figure 8A), characterized by strong upregulation of NF-κB pathway components (NFKB1, NFKB2, RELB), cytokines (IL6, IL1B, TNF), and chemokines (CXCL1, CXCL8, CCL2, CCL3, CCL4), as well as co-stimulatory molecules (CD80, ICOSLG) and matrix remodeling genes (FN1, THBS1, MMP9). This profile was accompanied by downregulation of metabolic and antioxidant regulators (FABP4, CAT, PDK2), indicating a shift toward an activated, inflammatory phenotype. GSEA analysis (Figure 8B) confirmed identified enrichment of pathways linked to inflammatory/immune response, NFκB nuclear import, response to LPS and cell proliferation, consistent with an activated phenotype. Notably, co-treatment with FL-ANGPTL3 counteracted FNIII-induced activation, suppressing inflammatory mediators (TNF, IL6, IL1B, IL12B, IL23A, CD80, CCL2, CCL4, CCL18) while enhancing genes linked to repair and immune regulation (SPP1, PDPN, APOE, GPNMB), consistent with an anti-inflammatory and pro-resolution effect (Figure 8C and F). C-ANGPTL3 exhibited intermediate suppression (Figure 8C and E), whereas LNA043 showed minimal impact on FNIII-driven transcriptional changes (Figure 8C-D).

**Figure 8.**
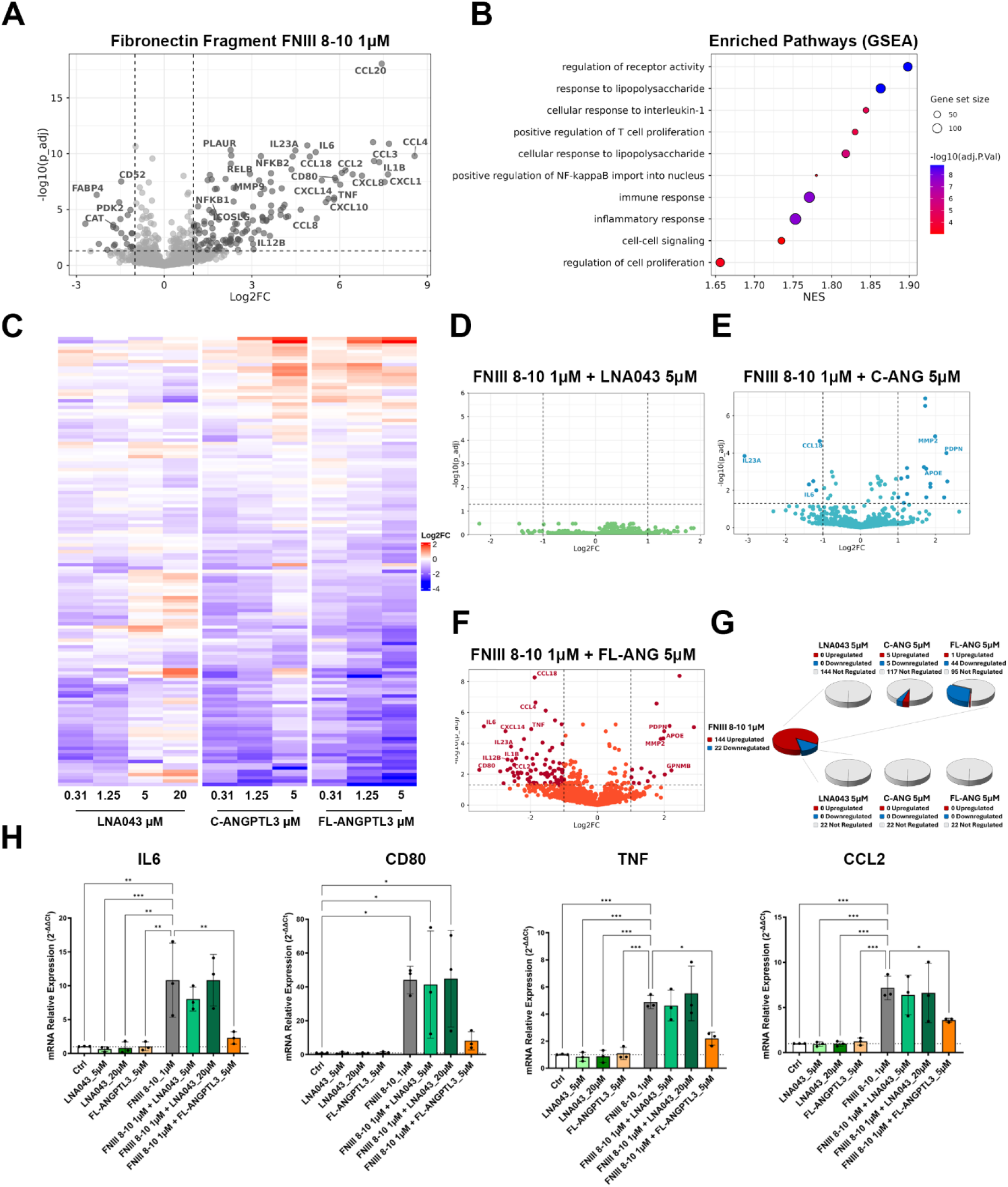
Targeted RASL-Seq transcriptomic analysis of THP-1 treated with FNIII 8-10. (A) Volcano plots illustrating differentially expressed genes (DEGs) after 24 hours treatment with 1 µM of FNIII 8-10. Data are expressed as log2 fold change (LOG2FC, x-axis) relative to untreated control, and significance is expressed as -log10 adjusted p-value (-log10(p_adj), y axis). Genes are considered significantly regulated above thresholds of LOG2FC ≥ 1 or ≤ –1 and –log10(p_adj) ≥ 1.3 (p_adj < 0.05). (B) Bubble plot showing the pathway enrichment analysis (GSEA). Pathways significantly enriched in FNIII 8-10 treated THP-1 are shown, with the x-axis representing the normalized enrichment score (NES). The size of the bubbles indicates the number of genes (Gene set size), and the color intensity indicates statistical significance (-log10(p_adj)). (C) Heatmap showing how the response of the genes significantly regulated by 1 µM FNIII 8-10 alone in panel (A) changes after 24 hours treatment with increasing concentrations of LNA043, C-ANGPTL3, or FL-ANGPTL3 in the presence of 1 µM FNIII 8-10. Genes are clustered by hierarchical analysis of log₂ fold-change values. Color scale represents relative expression changes, with red indicating upregulation and blue indicating downregulation. (D, E, F) Volcano plots illustrating DEGs after 24 hours treatment with 5 µM of LNA043 (D), C-ANGPTL3 (E), FL-ANGPTL3 (F) in the presence of 1 µM FNIII 8-10. Data are expressed as log2 fold change (LOG2FC, x-axis) relative to 1 µM FNIII 8-10. (G) Pie charts showing the number of statistically significant DEGs induced by 1 µM FNIII 8-10 (144 upregulated, 22 downregulated) and the effect of each treatment condition. The corresponding pie charts display how many of the FNIII 8–10–regulated genes are further upregulated (red), downregulated (blue), or remain unaffected (grey). (H) qPCR validation of top representative genes after treatment with LNA043 (5, 20 µM), FL-ANGPTL3 (5 µM). Data are shown as mean ± SD from 3 independent experiments. *p < 0.05, **p < 0.01, ***p < 0.001.

Importantly, among the 140 genes significantly upregulated by FNIII, FL-ANGPTL3 reduced the expression of 44, while C-ANGPTL3 affected only 5. Furthermore, neither C-ANGPTL3 nor FL-ANGPTL3 counter-regulated any of the 22 genes suppressed by FNIII (Figure 8G). We confirmed the RASL-Seq findings by qPCR on key inflammation-related genes regulated by FL-ANGPTL3 after 24 hours of treatment with LNA043 (5, 20 µM) or FL-ANGPTL3 (5 µM) in the presence of 1 µM FNIII 8-10. In line with the transcriptomic data, FL-ANGPTL3 markedly reduced FNIII 8–10–induced upregulation of IL6, CD80, TNF, and CCL2. Conversely, LNA043 did not counteract these changes (Figure 8H).

## Discussion

The primary aim of this study was to elucidate the mechanism of action of LNA043 in the context of ECM–integrin signaling within OA-relevant stromal and immune cells, while simultaneously deepening our understanding of ANGPTL3 biology by comparing LNA043 to its parental full-length and C-terminal FBN domain. Here, we show that FL-ANGPTL3, C-ANGPTL3 and LNA043 co-localize with fibronectin and α5β1 integrin at fibrillar adhesions, interact with FNIII 12–14 and engage α5β1 integrin, indicating that a truncated C-terminal domain is sufficient for these interactions. AlphaFold modeling supports the formation of a ternary complex between FN,α5β1 integrin, and all three proteins. C-ANGPTL3 largely recapitulated FL-ANGPTL3 activity, including ihMSCs adhesion, monocyte migration, proliferation, and transcriptomic modulation, whereas a further truncation of additional 17aa in LNA043 markedly attenuated these responses. While FL- and C-ANGPTL3 induced robust β1 integrin-dependent ihMSC adhesion and counter-regulated FNIII 8–10–driven inflammatory programs in monocytes, LNA043 selectively promoted β1-dependent monocyte migration without supporting ihMSC adhesion, THP-1 proliferation, or inflammatory gene suppression, highlighting its distinct and restricted functional profile. Together, these data suggest that both LNA043 and C-ANGPTL3 act via β1 integrins but have lower intrinsic activity compared to FL-ANGPTL3. Consistent with these functional differences, β1 integrin blockade completely abolished C-ANGPTL3–mediated adhesion, whereas FL-ANGPTL3–dependent adhesion was only partially inhibited and became significant only at lower coating concentrations. This suggests that β1 integrins are required but not solely sufficient to account for the full activity of FL-ANGPTL3. In contrast, THP-1 migration responses to all three proteins were only partially inhibited, which may reflect either additional contributing mechanisms or incomplete β1 integrin blockade under the experimental conditions or antibody concentrations used.

The reduced activity of LNA043 may be related to structural elements that were lost upon truncation. The 17aa missing in LNA043 may include motifs or co-factor sites that, while not required for integrin recognition, could improve interaction stability or multivalency in FL- and C-ANGPTL3. In addition, the stronger effect of C-ANGPTL3 compared to LNA043 suggests the presence of elements within the longer C-terminal fragment that could help maintain a more functional integrin-binding surface. An additional factor that may contribute to the reduced activity of LNA043 is the absence of O-glycosylation motifs that are present in C-ANGPTL3. ANGPTL3 undergoes site-specific O-glycosylation close to its pro-protein convertase cleavage site, with two GalNAc O-glycosylation sites mapped to the TT226 motif in the C-terminal region (43,44). These O-glycans were shown to modulate ANGPTL3 processing and conformation, and could influence accessibility of the C-terminal FBN domain by altering structural stability and protease interactions. Although O-glycosylation can, in principle, modulate protein folding, receptor-binding geometry, and multivalent presentation (45,46), there is currently no evidence that O-glycans present on ANGPTL3 regulate cell adhesion, integrin binding, or downstream signaling. Nonetheless, the O-glycosylation site located immediately C-terminal to the ANGPTL3 pro-protein convertase cleavage region represents a structural element that is absent in LNA043. While speculative, the absence of these O-glycan structures in LNA043 may reduce its ability to maintain an optimal integrin-binding interface or activation. Targeted mutagenesis of the known O-glycosylation sites in C-ANGPTL3, glyco-engineering of LNA043, and comparative biophysical measurements could help determine whether the absence of these glycans contributes to the reduced LNA043 bioactivity observed in the analyzed in vitro assays. Importantly, we can exclude the possibility that the reduced activity of LNA043 is caused by the K423Q point mutation introduced to prevent LNA043 proteolysis in the production cell line. As shown in the adhesion and migration assays, neither reverting the mutation nor expressing LNA043 in the same production system used for FL- and C-ANGPTL3 restored activity, demonstrating that the mutation or potential differences in glycosylation do not affect integrin-dependent cell adhesion or migration.

While ANGPTL3 has been studied primarily for its regulation of lipid metabolism via its N-terminal domain, the function of the C-terminal FBN-like domain remains less defined. Camenisch et al. showed that this domain supports endothelial cell adhesion and migration via integrin αVβ3 and is sufficient to promote angiogenesis in vivo (25). Our findings extend these observations by showing that the ANGPTL3 FBN-like domain also binds fibronectin and engages integrin α5β1, revealing an additional α5β1-dependent function in ihMSCs and monocyte. Consistent with this, other ANGPTL family members regulate cartilage and ECM biology through integrin-dependent mechanisms: ANGPTL2 promotes chondrogenesis but drives OA-associated inflammation via α5β1-mediated NF-κB activation (31,47), while ANGPTL4 has different roles in cartilage and it associates with fibronectin through its FBN-like domain and modulates cell–matrix signaling during tissue repair (48–50). Together, these findings suggest that fibronectin binding by ANGPTL FBN-like domains represents a conserved mechanism for regulating integrin-dependent signaling.

Moreover, through targeted RASL-Seq profiling our study in human monocytes identified a previously unrecognized immunomodulatory role of ANGPTL3. This approach revealed that FL-ANGPTL3 and C-ANGPTL3 induce a broad transcriptional reprogramming, upregulating the expression of chemokines, ECM-remodeling factors and regulators of myeloid activation, while LNA043 shows minimal transcriptomic activity. Importantly, the immunomodulatory capacity was not limited to simple stimulation. By mimicking the inflammatory environment of OA using the FNIII 8–10 fragment we discovered that FL-ANGPTL3 counter-regulates FNIII-8-10-induced inflammatory transcription, suppressing core NFκB-driven mediators such as IL1B, IL6, TNF, IL12B, IL23A, while enhancing transcripts associated with tissue repair and resolution. C-ANGPTL3 displayed a similar but less pronounced effect, while LNA043 lacked counter-regulatory capacity altogether. These findings indicate that ANGPTL3 is a modulator of immune activation capable of counterbalancing matrix-derived fibronectin fragments signaling in monocytes.

From a translational point of view, the reduced activity of LNA043 in immune cells compared to ANGPTL3 is well aligned with its established safety and tolerability profile. In preclinical cartilage-injury and OA models, LNA043 regenerated hyaline-like cartilage without inducing angiogenesis or inflammatory activation, indicating that it retains cartilage repair activity without promoting immune responses and vascular remodeling. These observations were reinforced in the first-in-human randomized phase 1 trial, where intra-articular LNA043 showed excellent tolerability, no adverse effects, no immunogenicity, and clear penetration into human cartilage (16). In this context, the ability of LNA043 to bind fibronectin and localize to fibrillar adhesions suggests that its activity is spatially restricted within fibronectin-rich extracellular matrix, potentially enabling sustained local repair signaling while limiting immune activation. Together, these data support a model in which LNA043 acts as a precisely tuned, ECM-anchored, cartilage repair protein.

Some limitations should be acknowledged. While structural modeling and PLA data support the formation of ternary complexes, direct biophysical validation is still required, for example through SPR-based binding analyses or high-resolution structural approaches such as cryo-EM. In addition, all functional data were generated in vitro, and it remains to be determined whether the immunomodulatory effects of FL-ANGPTL3 and C-ANGPTL3, including suppression of FNIII 8–10–driven inflammatory responses, are recapitulated in vivo and whether the distinct activity profile of LNA043 translates into a more selective or safer immunomodulatory response. Finally, transcriptomic analyses were limited to a targeted RASL-Seq panel, and broader, unbiased profiling in primary human cells or ex vivo joint models will be important to further refine the mechanisms of action of ANGPTL3 and LNA043.

Together, our findings showed a novel involvement of ANGPTL3 in the fibronectin–α5β1 signaling pathway and establish a first mechanistic explanation for the divergent biological and therapeutic properties of LNA043 compared to native ANGPTL3. These data provide a rationale for the continued development of LNA043 as therapy for OA and cartilage injury repair. Moreover, the discovery that ANGPTL3 can co-engage fibronectin and α5β1, modulate inflammatory transcriptional programs, and counteract matrix-derived DAMP signaling in monocytes suggests that ANGPTL3-derived proteins, or domain-specific variants, could hold therapeutic potential in inflammatory conditions in which integrin- and ECM-mediated immune activation play central roles.

## Acknowledgment

This research was supported by the Novartis Postdoctoral Fellowship Program, Biomedical Education and Innovation, Novartis Biomedical Research, Novartis Pharma AG. The content is solely the responsibility of the authors.

## Author contributions

Study concept and design: A.G, C.H., M.F. Data acquisition and analysis: A.G., S.C., E.G., T.L., T.H., G.K., F.K. Protein production: K.N., J.S., M.V. Drafting of the manuscript: A.G., C.H., M.F. Revision of the manuscript: A.G., S.C., E.G., T.L., T.H., G.K., F.K., K.N., J.S., M.V., N.G., C.H., M.F. Approval of the final manuscript: all authors.

## Methods

### Cell culture

The immortalized human bone marrow-derived mesenchymal stem cells (ihMSC) UE7T-13 (JCRB Cell Bank) were cultured and expanded in Mesenchymal Stem Cell Growth Medium (MSCGM®) supplemented with MSCGM® SingleQuots® Supplement Kit (Lonza). Immortalized hMSC were cultured in 5% CO₂ in a humidified incubator at 37°C until they reached 80-90% confluence. THP-1 (ATCC) were cultured in RPMI-1640 medium (ATCC modification) (Gibco) supplemented with 10% fetal bovine serum (FBS), 1% penicillin-streptomycin and 50 μM β-mercaptoethanol (Gibco). Cells were kept at 37°C in a humidified incubator with 5% CO₂ and were passaged upon reaching a concentration of 1 x 10⁶ cells/ml.

### CRISPR/CasG Knockout of ITGA5, ITGB3, and FN1 in ihMSCs

Stable knockout (KO) cell lines were generated in ihMSCs UE7T-13 (JCRB Cell Bank) using CRISPR/Cas9 ribonucleoprotein (RNP) complexes delivered via electroporation with the Neon® Transfection System (MPK5000, Invitrogen). For each gene of interest, two guide RNAs (gRNAs) were designed using the CRISPOR tool (http://crispor.tefor.net/), selecting sequences with high specificity and predicted editing efficiency. The gRNA sequences used were as follows: integrin α5 (ITGA5), 5′-CGGGGGCTTCAACTTAGACG-3′ and 5′-TTTACCGGCCGGGAACAGAC-3′; fibronectin (FN1), 5′-AGCTGCACGAACATCGGTGA-3′ and 5′-CAAATGCTTTGCACGTGCCT-3′; and integrin β3 (ITGB3), 5′-AGGCGGACGAGATGCGAGCG-3′ and 5′-CTGGCGGGCGTTGGCGTAGG-3′. A 22 bp double-stranded multi-stop codon (MSC) oligonucleotide (5′-TAA CTA GTT GAT TAT TTA GTC A-3′; PAGE purified, Microsynth, Balgach, Switzerland) was included to enhance knockout efficiency. Alt-R® CRISPR-Cas9 tracrRNA and crRNA (Integrated DNA Technologies, IDT, Coralville, Iowa, USA) were annealed in Nuclease-Free Duplex Buffer (IDT) at 95°C for 5 minutes and cooled to room temperature. The gRNA duplex was complexed with Alt-R® S.p. HiFi Cas9 Nuclease V3 (IDT) and incubated for 10 minutes at room temperature. Cells were prepared by trypsinization, washed, and resuspended in Buffer R at a final concentration of 2 × 10⁶ cells per 100 μL. The RNP complex, MSC oligonucleotide, and cells were combined and briefly incubated prior to electroporation. Electroporation was performed using 10 μL Neon® tips with cell line-specific parameters. Following electroporation, cells were transferred to antibiotic-free growth medium and seeded in appropriate vessels. One day post-transfection, the medium was replaced with growth medium containing penicillin/streptomycin and refreshed every 2–3 days. Cells were expanded and harvested for downstream analyses including genotyping, protein expression profiling, and functional assays. CRISPR/Cas9 editing efficiency for integrin targets was assessed by flow cytometry, and positively edited cells were sorted using a BD Fortessa cell sorter (BD Life Sciences). Knockout validation was performed by Western blot and quantitative PCR (qPCR) (data not shown). For FN1, knockout efficiency was evaluated by qPCR and Western blot (data not shown).

### Recombinant Production and Purification of LNA043, C-ANGPTL3, FL-ANGPTL3, and Fibronectin fragments

LNA043, a recombinant C-terminal ANGPTL3-derived protein generated by truncations and mutagenesis, was produced in Chinese hamster ovary (CHO) cells (Novartis Pharma AG) as previously described (16). Coding sequences corresponding to full-length human ANGPTL3 (FL-ANGPTL3; aa 1-460) and to the C-terminal fibrinogen-like domain (C-ANGPTL3; aa 225–460) were cloned into expression vectors alongside codon-optimized constructs encoding LNA043 variants (wild-type or K423Q) that were used for comparative analysis. For all constructs, a C-terminal His-tag sequence was included to enable affinity purification. All three proteins were produced using identical Expi293 (Expi293F™, Thermo Fisher Scientific) expression conditions. Transient transfections were performed at a cell density of 3.0 × 10⁶ cells/ml, and cultures received transfection enhancer 24 hours later before transfer to an Infors HT Multitron Triple Incubator Shaker operating at 37 °C, 5% CO₂, and 120 rpm. Five days post-transfection, conditioned medium was harvested, clarified by centrifugation, filtered through a 0.22 µm membrane, and supplemented with 1 mM NiCl₂. For purification, clarified supernatants were loaded onto an AKTA Pure chromatography system fitted with two Qiagen Ni-NTA Superflow columns in series. Columns were equilibrated in 20 mM Tris-HCl pH 8.0 and 150 mM NaCl, then washed with 20 column volumes (CV) of equilibration buffer followed by 10 CV of wash buffer containing 10 mM imidazole. All washing steps were performed using a peristaltic pump at a flow rate exceeding 0.5 ml/min, consistent with the operational parameters shown in the purification workflow. Bound proteins were eluted using 20 mM Tris-HCl pH 8.0, 150 mM NaCl, and 250 mM imidazole. Eluted fractions were pooled and buffer-exchanged into 20 mM Tris-HCl pH 7.5 containing 150 mM NaCl and concentrated before freezing for downstream applications.

Coding sequences for Fibronectin type-III domains 8-10 (aa 1267-1540) and Heparin-binding 2 domain (aa 1812-2084) of human Fibronectin (UniProt ID: P02751) were subcloned into the pET100D expression vector, introducing an N-terminal His6-tag followed by a TEV protease cleavage site. The resulting plasmids were transformed into E. coli BL21 Gold (DE3) cells for protein production. For expression, cultures were grown and induced according to standard procedures, followed by incubation for 14 hours at 20 °C. Cells were harvested and lysed, and clarified lysates were subjected to immobilized metal affinity chromatography using a HisTrap Excel column. Following elution, samples were incubated with TEV protease to remove the N-terminal His6-tag. Proteins were subsequently polished by size-exclusion chromatography using a Superdex 75 26/600 column, and peak fractions corresponding to the target proteins were collected for downstream applications.

### Immunofluorescence staining

ihMSCs were seeded into μ-Plate 96 Well Black ibiTreat plates (ibidi GmbH,). After incubation with LNA043, C-, or FL-ANGPTL3 for 1 hour at 37°C, cells were fixed with 4% paraformaldehyde and prepared for immunostaining or proximity ligation assay (PLA). Immunofluorescence staining was performed with the following primary antibodies and dilutions: mouse anti-ANGPTL3 (clone OTI5E6, LSBio) at 1:200; mouse anti-integrin α5β1(clone HA5, Sigma-Aldrich) at a dilution of 1:100; mouse anti-fibronectin (clone IST9, Abcam) at 1:200; mouse anti-integrin αVβ3 (clone LM609, Merck KGaA) at 1:200. Primary antibodies were incubated either overnight at 4°C or for one hour at room temperature. Subsequently, Alexa-Fluor secondary antibodies (Invitrogen) were applied at a dilution of 1:400 for one hour at room temperature. Nuclei were counterstained with DAPI. Fluorescence images were acquired with an Olympus FV3000 confocal laser scanning microscope (Olympus).

### ELISA for protein binding to fibronectin and fibronectin fragments

ELISA plates (Nunc® Maxisorp™; Nunc) were coated overnight at 4°C with 50 nM fibronectin, fibronectin fragments, LNA043, or C- and FL-ANGPTL3 in PBS. Coating proteins included full-length human plasma fibronectin (Sigma), 70 kDa fibronectin fragment (Sigma), hsFN8-10 (FNIII 8-10, aa 1267–1540), hsFN heparin-binding domain (FNIII 12-14 HBDII, aa 1812–2084), LNA043, C- and FL-ANGPTL3 (Novartis). Wells were washed twice with PBS containing 0.05% Tween-20 (PBS-T) and blocked with 2% (wt/vol) BSA in PBS-T for 1 hour at room temperature. After washing, wells were incubated with increasing doses of LNA043, C-, FL-ANGPTL3, full-length human plasma fibronectin or vehicle control diluted in PBS-T containing 0.1% BSA for 1 hour at 37°C. Plates were washed three times with PBS-T and incubated with biotinylated anti-ANGPTL3 antibody (RCD Systems) at 0.25 μg/ml or biotinylated anti-Fibronectin antibody (Rockland) at 0.1 μg/ml for 1 hour at room temperature. After three washes, wells were incubated with streptavidin-HRP conjugate (RCD Systems) for 1 hour. Detection was performed using Tetramethylbenzidine substrate (TMB, RCD Systems) for 20 minutes at room temperature, and the reaction was stopped with 2N H₂SO₄. Absorbance was measured at 450 nm and 540 nm using a microplate reader (Agilent Technologies).

### In silico structure predictions with AlphaFold

The structures of the complexes were predicted by AlphaFold 2.3.2 (51) using the docker in multimer mode and employing the full structural repository (max_template_date disabled). The primary sequence of LNA043 was previously disclosed (16), while the sequence of FL-ANGPTL3 was retrieved from NCBI NLM (RefSeq: NP_055310.1). Sequences for integrin subunits α5 and β1 were extracted from PDB entry 7NWL, and those of αV, α4 and fibronectin were obtained from UniProt IDs P06756-1, P13612-1 and P02751-15, respectively. The headpieces regions of integrins and the HBDII domain of fibronectin were manually extracted for modeling. For each complex, 25 conformers were generated and further analyzed using PyMOL v3.1.6. All conformers were aligned on the ANGPTL3 molecule and manually inspected and classified into recurring similar conformations. Figures and video were generated using PyMOL, based on the best predicted model for each conformation, which was subsequently used as the starting structure for molecular dynamics (MD) simulations.

### Molecular dynamics simulations

MD simulations were performed using GROMACS version 2024.1 (DOI 10.5281/zenodo.10090355). Proteins were modelled using the OPLS/AA force field, and water molecules were modelled as 3-site rigid with partial charge assignment (SPC/E model). Each protein complex was placed in a periodic boundary condition cubic simulation box maintaining a minimum distance l of 2.0 nm from the protein to the box edge. The system was solvated with water and neutralized with Na^+^ and Cl^-^ ions to a target concentration of 100 mM. Energy minimization was conducted using the steepest descent algorithm over 50,000 steps with a step size of 0.01 nm. Long-range electrostatic interactions were treated using the Particle Mesh Ewald (PME) method. For production runs, the Verlet leapfrog algorithm with a 2 fs time step was used. Short-range interactions were calculated up to a cutoff of 1.4 nm. The long-range Coulombic interactions were estimated using the PME method with a cutoff distance of 1.4 nm. The production run was performed in NPT ensemble at 1 bar and 300 K. The stochastic velocity and pressure rescaling thermostat and barostat were used with frictions of 1.0 and 2.0 ps for temperature or pressure coupling, respectively Each simulation ran for 100 ns, generating 10,000 independent conformations. The first 20 ns were discarded to allow for equilibration, which was validated by monitoring fluctuations in total energy and system density. The Coulombic and Lenard-Jones interacting energies were compiled between two binding interfaces: LNA043 and the binding partner (either integrin α5β1, considered as a single binder, or Fibronectin’s HBDII domain). For C-and FL-ANGPTL3, the binding interface was limited to the FBN-like domain (i.e., LNA043 sequence) to ensure comparability. Average interaction energies and RMSD values are reported in Supplementary Table.

### Proximity Ligation Assay (PLA)

Protein-protein interactions were analyzed using the Duolink® PLA Fluorescence kit (Sigma-Aldrich) following the manufacturer’s protocol. Briefly, 5 x 10^3^ ihMSCs were seeded into μ-Plate 96 Well Black ibiTreat plates (ibidi GmbH). After overnight incubation, cells were incubated with 300 µg/ml (12 µM) of LNA043 for 1 hour at 37°C and fixed with 4% paraformaldehyde. Blocking was performed using BSA 2% and FBS 5% in PBS, followed by incubation with primary antibody pairs: rabbit anti-ANGPTL3 (Rockland) at 1:100 with either mouse anti-Integrin α5β1 (clone HA5) or mouse anti-Fibronectin (clone IST9) both at 1:200. Species-specific PLA probes (PLUS and MINUS) were applied, and ligation and amplification steps were performed to generate discrete fluorescent signals indicating protein proximity (<40 nm). Nuclei were counterstained with DAPI, and PLA signals were visualized using an Olympus FV3000 confocal laser scanning microscope (Olympus).

### Assessment of integrin-mediated adhesion of mesenchymal stem cells to ANGTPL3 domains

96-well plates (Nunc® Maxisorp™; Nunc) were coated overnight at 4°C with different concentrations of LNA043, C and FL-ANGPTL3. Uncoated wells served as BSA negative controls. Wells were washed twice with PBS without Ca²⁺ and Mg²⁺ (Gibco) and blocked with 2% (wt/vol) BSA in PBS for 1 hour at room temperature. ihMSC were detached using Accutase™ (STEMCELL technologies), neutralized with growth medium (MSCGM BulletKit; Lonza), and washed twice with serum-free DMEM (Gibco). Cells were resuspended in adhesion buffer (DMEM + 1 mM CaCl₂ + 1 mM MgCl₂ + 0.5 mM MnCl₂) and seeded at 40,000 cells per well in 100μl. For integrin blocking experiments, cells were pre-incubated for 30 minutes at 37°C with integrin-blocking antibodies at 20 µg/mL, including anti-α5β1 (clone JBS5), anti-β1 (clone mAb13), and anti-αVβ3 (clone LM609) (Merck KGaA); anti-α4 (clone 2B4, RCD Systems); isotype control (Thermo Fisher Scientific) or 5 nM EDTA (Sigma-Aldrich). Following 1 hour incubation at 37°C, the wells were carefully washed with PBS to eliminate non-adherent cells. Subsequently, the adherent cells were fixed using 4% paraformaldehyde (Electron Microscopy Sciences) for 10 minutes at room temperature. Fixed cells were stained with 0.5% crystal violet (Sigma-Aldrich) for 10 minutes, washed with deionized water, and air-dried. Crystal violet was solubilized with 150 µL of pure methanol (VWR) for 20 minutes with gentle agitation. Absorbance was measured at 570 nm using a microplate reader (Agilent Technologies).

### Peripheral Blood Mononuclear Cell (PBMC) and human monocytes isolation

Primary human blood cells were obtained from healthy adult donors through the Novartis employee donor program at Novartis Pharma AG (Basel, Switzerland), following written informed consent. The donor program and experimental protocols involving human samples were reviewed and approved by the Novartis BioSample Compliance function under the oversight of Ethics, Risk & Compliance (ERC), in accordance with institutional ethical standards, the Declaration of Helsinki, and local regulatory requirements. The donation program consequently obtained ethics approval for its operations, granted by the Ethikkommission Nordwest-und Zentralschweiz (EKNZ). Samples were fully anonymized prior to analysis. PBMCs were isolated from fresh human blood using LeucoSep™ tubes (Greiner Bio-One GmbH) pre-filled with Ficoll-Paque™ PLUS (Cytiva). A total of 15 mL of Ficoll-Paque was added to each of LeucoSep tubes. To settle the porous barrier and eliminate air bubbles, tubes were centrifuged at 1000 × g for 30 seconds at room temperature. Blood was diluted 1:1 with sterile PBS and carefully added to the LeucoSep tube. Samples were centrifuged at 800 × g for 15 minutes at room temperature with the break off. The mononuclear cell layer was collected from the interface, transferred to a new tube, and washed twice with PBS by centrifugation at 300 × g for 10 minutes to remove residual Ficoll and platelets. Cell viability and concentration were assessed using trypan blue exclusion and an automated cell counter. Isolated PBMCs were used immediately for downstream experiments. CD14^+^CD16^−^ monocytes were subsequently isolated from PBMCs (three different donors) using the Classical Human Monocyte Isolation Kit (Miltenyi Biotec), following the manufacturer’s protocol. Briefly, non-monocyte populations including T cells, B cells, NK cells, dendritic cells, and basophils were indirectly magnetically labeled using a cocktail of biotin-conjugated antibodies and Anti-Biotin MicroBeads. Labeled cells were depleted using LS Columns and a MidiMACS™ Separator, yielding a highly pure population of untouched classical monocytes. Isolated monocytes were used immediately for downstream applications.

### Monocytes migration assay

Migration assays were performed using 24-well plates (Corning) fitted with 6.5 mm Transwell® inserts containing 5.0 µm pore polycarbonate membranes (Corning Inc). Inserts were placed into sterile, non-treated 24-well clear plates (Costar®, Corning Inc.). THP-1 cells were resuspended and counted, then washed twice and adjusted to the desired concentration with serum-free RPMI medium. A total of 0.25 x 10⁶ cells in 100 µL were seeded into the upper chamber of each Transwell insert. The lower chamber was filled with 600 µL of migration medium containing either vehicle control, LNA043, C-ANGPTL3 or FL-ANGPTL3. Cells were allowed to migrate for 6 and 24 hours at 37°C. After incubation, inserts were transferred to empty wells, and the contents of the lower chamber were collected, centrifuged at 300 × g for 5 minutes, and resuspended in a smaller volume of serum-free RPMI medium. Migrated cells were quantified using a Countess™ automated cell counter (Thermo Fisher Scientific). To evaluate integrin dependency in THP-1 cell migration, cells were pre-incubated with integrin-blocking antibodies at 20 µg/mL for 30 minutes at 37°C prior to seeding into Transwell® inserts. The following antibodies were used: anti-α5β1 (clone JBS5), anti-β1 (clone mAb13), and anti-αVβ3 (clone LM609) (all from Merck KGaA); anti-α4 (clone 2B4, RCD Systems); and mouse IgG isotype control (Thermo Fisher Scientific). For migration assays using primary human monocytes isolated with Classical Human Monocyte Isolation Kit (Miltenyi Biotec GmbH) as described above, 0.15 x 10⁶ cells per Transwell insert were seeded in 100 µL of serum-free (SF) RPMI medium.

### Assessment of THP-1 cell proliferation

THP-1 cells were seeded at a density of 20,000 cells per well in a 96-well plate (Corning) and treated with LNA043, C-ANGPTL3, or FL-ANGPTL3 for 24 hours in SF RPMI under standard culture conditions (37 °C, 5% CO₂). Cell proliferation was quantified using the CyQUANT™ Direct Cell Proliferation Assay Kit (Thermo Fisher Scientific) according to the manufacturer’s instructions. Briefly, after treatment, CyQUANT™ Direct reagent was added directly to the wells and incubated for 60 min at 37 °C. Fluorescence intensity was measured using a microplate reader (excitation/emission: 480/535 nm), and cell number was calculated based on a standard curve generated from known THP-1 cell densities.

### Flow cytometry analysis of surface integrin expression

THP-1 cells and primary human monocytes were stained for surface integrin expression using fluorophore-conjugated antibodies. Cells were first incubated with Human TruStain FcX™ (BioLegend) to block Fc receptors and reduce non-specific binding. Following Fc blocking, cells were stained with PE-conjugated anti-α5β1 (mouse IgG2b, clone 05; Sino Biological) and APC-conjugated anti-αVβ3 (mouse IgG1, clone 23C6; BioLegend). Corresponding isotype controls were included: PE-conjugated mouse IgG2b κ and APC-conjugated mouse IgG1 κ (BioLegend). Staining was performed in FACS buffer (PBS + 2% FBS) for 30 minutes at 4°C in the dark. After staining, cells were washed twice with cold FACS buffer and resuspended for acquisition on a CytoFLEX S flow cytometer (Beckman Coulter). Data were analyzed using FlowJo™ Software v10 (BD Life Sciences).

### THP-1 treatment for gene expression profiling and protein quantification

THP-1 cells were cultured under standard culture conditions until reaching a density of 1 million cells/ml. For RASL-Seq profiling, cells were harvested and washed with serum-free medium to remove residual serum, followed by a 1 hour starvation period. Subsequently, cells were seeded into 384-well plates (Greiner Bio-One) at 20,000 cells per well and treated with increasing concentrations of LNA043, C-ANGPTL3, or FL-ANGPTL3 (n=6). Incubation continued for 24 hours. For FNIII 8-10 treatment, 1 μM of FNIII 8-10 was added after the initial 1 hour LNA043 or ANGPTL3 treatment, and cells were further incubated for 24 hours. After incubation, plates were centrifuged at 200 × g for 5 minutes to pellet the cells. Supernatants were removed, and cells were lysed using a Proteinase K solution (Thermo Fisher Scientific) in PK Buffer for MagMAX™ (Life Technologies) at a 1:24 ratio. Lysates were stored at −80 °C and submitted for RASL-Seq processing. For qPCR validation, 0.2 million THP-1 cells were seeded into 96-well plates (Corning Inc.) and treated as described for RASL-Seq. After 24 hours, cells were pelleted and lysed in TRIzol™ Reagent (Thermo Fisher Scientific). Supernatants were centrifuged at 14,000 x g for 10 minutes to remove cellular debris and subsequently stored at -80°C pending analysis.

### RASL-Seq analysis

RASL-Seq screening was conducted in accordance with the probe design principles and experimental workflow described by Li et al. (52). Using this methodology, transcript abundance for 1,764 genes was quantified. The gene panel comprised biomarkers relevant to myeloid biology as well as genes associated with general cellular processes, selected based on both internal and external datasets. For most targets, two to three independent RASL-Seq probe sets were employed to ensure robustness. Count matrix was obtained by mapping the raw reads to the probe sequences, and samples with <15,000 mapped reads were removed. Read counts were then normalized to Reads Per Million (RPM) based on total mapped reads. For genes represented by three independent probe sets, the probe showing the strongest and most consistent expression, benchmarking against an internal RNA-Seq dataset (not shown), was selected as the representative measure. Consistency was assessed using inter-replicate Pearson correlation of log₂(RPM + 1) values across all genes. Quality filtering excluded samples with a maximum inter-replicate correlation < 0.8 or with > 40% undetected probes relative to the best-performing replicate. To address technical variability, batch effects across plates were corrected using pyCombat, followed by quantile normalization of log₂(RPM + 1) values to harmonize expression distributions across replicates. Differential gene expression analyses were subsequently performed using the limma framework in R (53).

Gene set enrichment analysis (GSEA) was performed using the *fgsea* R package version 1.34.0 (54). Genes were ranked by log₂ fold change from differential expression analysis, and enrichment was assessed using GO gene sets with the fgsea function under default parameters. Normalized enrichment scores (NES) and Benjamini–Hochberg adjusted p-values were used for downstream analyses.

### Quantitative real-time PCR

THP-1 cells were lysed in TRIzol™ Reagent (Thermo Fisher Scientific), and total RNA was extracted using the RiboPure™ RNA 96-well purification kit (Zymo Research Europe GmbH) according to the manufacturer’s instructions. RNA concentration and purity were assessed spectrophotometrically, and equal amounts of RNA were reverse transcribed into cDNA using the SuperScript III Reverse Transcriptase (Thermo Fisher Scientific). Quantitative real-time PCR (qRT-PCR) was performed on a7900HT Real-Time PCR System (Applied Biosystems) using human-specific TaqMan gene expression assays (Thermo Fisher Scientific). Reactions were run in duplicate and normalized to GAPDH expression. Relative mRNA levels were calculated using the 2^−ΔΔCt method. Expression of genes of interest was determined using the following human specific TaqMan gene expression assays (Thermo Fisher Scientific): GAPDH (Hs99999905_m1), AXL (Hs01064444_m1), CCL2 (Hs00234140_m1), CCL20 (Hs00355476_m1), CD80 (Hs01045161_m1), FABP4 (Hs01086177_m1), FN1 (Hs01549976_m1), IL1B (Hs01555410_m1), IL6 (Hs00174131_m1), MMP2 (Hs01548727_m1), PDPN (Hs00366766_m1), SKIL (Hs00180524_m1), TNF (Hs00174128_m1).

### ELISA for protein quantification in THP-1 supernatant

The concentrations of FN1, CCL20, and MMP2 were measured using the U-PLEX electrochemiluminescence immunoassay platform (Meso Scale Discovery) following the manufacturer’s protocol. Briefly, U-PLEX 96-well plates were coated with linker-coupled capture antibodies for the three analytes and subsequently incubated with standards and appropriately diluted samples. After washing, Sulfo-tag-labeled detection antibodies were added and incubated. Following additional washes, MSD Read Buffer was dispensed into each well, and plates were immediately analyzed using the Meso Sector 600MM instrument. Electrochemiluminescent signals were quantified using MSD Discovery Workbench (Meso Scale Discovery) and concentrations were expressed in pg/ml.

### Statistics

Data are presented as mean ± standard deviation. Statistical analyses were performed using GraphPad Prism 10 software (GraphPad Software). Normality of data distribution was assessed prior to statistical testing. Depending on the outcome, multiple group comparisons were conducted using either parametric one-way analysis of variance (ANOVA) followed by Bonferroni post hoc test, or nonparametric Kruskal–Wallis test followed by Dunn’s post hoc test. Differences were considered statistically significant at p < 0.05.

## Data availability

The data supporting the findings of this study are available from the corresponding author upon reasonable request and subject to Novartis data-sharing policies.

## References

1. Tang S, Zhang C, Oo WM, Fu K, Risberg MA, Bierma-Zeinstra SM, et al. Osteoarthritis. Nat Rev Dis Primers. 2025 Dec 1;11(1). doi:10.1038/s41572-025-00594-6 PubMed PMID: 39948092.

2. Collins KH, Haugen IK, Neogi T, Guilak F. Osteoarthritis as a systemic disease. Nat Rev Rheumatol. 2025 Dec 3. doi:10.1038/s41584-025-01332-8

3. Yao Q, Wu X, Tao C, Gong W, Chen M, Qu M, et al. Osteoarthritis: pathogenic signaling pathways and therapeutic targets. Signal Transduction and Targeted Therapy. Springer Nature; 2023. doi:10.1038/s41392-023-01330-w PubMed PMID: 36737426.

4. Sanchez-Lopez E, Coras R, Torres A, Lane NE, Guma M. Synovial inflammation in osteoarthritis progression. Nature Reviews Rheumatology. Nature Research; 2022. p. 258–75. doi:10.1038/s41584-022-00749-9 PubMed PMID: 35165404.

5. Deng Z, Yang W, Zhao B, Yang Z, Li D, Yang F. Advances in research on M1/M2 macrophage polarization in the pathogenesis and treatment of osteoarthritis. Heliyon. Elsevier Ltd; 2025. doi:10.1016/j.heliyon.2025.e42881

6. Griffin T.M., Scanzello C.R. Innate inflammation and synovial macrophages. Experimental Rheumatology. 2019.

7. Haubruck P, Pinto MM, Moradi B, Little CB, Gentek R. Monocytes, Macrophages, and Their Potential Niches in Synovial Joints – Therapeutic Targets in Post-Traumatic Osteoarthritis? Frontiers in Immunology. Frontiers Media S.A.; 2021. doi:10.3389/fimmu.2021.763702 PubMed PMID: 34804052.

8. Lambert C, Zappia J, Sanchez C, Florin A, Dubuc JE, Henrotin Y. The Damage-Associated Molecular Patterns (DAMPs) as Potential Targets to Treat Osteoarthritis: Perspectives From a Review of the Literature. Frontiers in Medicine. Frontiers Media S.A.; 2021. doi:10.3389/fmed.2020.607186

9. Pérez-García S, Carrión M, Gutiérrez-Cañas I, Villanueva-Romero R, Castro D, Martínez C, et al. Profile of matrix-remodeling proteinases in osteoarthritis: Impact of fibronectin. Cells. MDPI; 2020. doi:10.3390/cells9010040 PubMed PMID: 31877874.

10. Reed KSM, Ulici V, Kim C, Chubinskaya S, Loeser RF, Phanstiel DH. Transcriptional response of human articular chondrocytes treated with fibronectin fragments: an in vitro model of the osteoarthritis phenotype. Osteoarthritis Cartilage. 2021 Feb 1;29(2):235–47. doi:10.1016/j.joca.2020.09.006 PubMed PMID: 33248223.

11. Barilla ML, Carsons SE. Fibronectin Fragments and Their Role in Inflammatory Arthritis. 2000.

12. Khan S, Cho H, Hasty KA, Brown T, Bhogoju S, Subramanian A. Synoviocyte– chondrocyte triculture model for early-stage PTOA: fibronectin fragment-induced catabolic effects in vitro and in vivo. Front Bioeng Biotechnol. 2025;13. doi:10.3389/fbioe.2025.1683333

13. Kragstrup T, Sohn D, Lepus C, Onuma K, Wang Q, Robinson W, et al. Tissue growth factor β stimulates fibroblast-like synovial cells to produce extra domain A fibronectin in osteoarthritis [Internet]. 2018. Available from: http://biorxiv.org/lookup/doi/10.1101/335562 doi:10.1101/335562

14. Miao MZ, Lee JS, Yamada KM, Loeser RF. Integrin signalling in joint development, homeostasis and osteoarthritis. Nature Reviews Rheumatology. Nature Research; 2024. p. 492–509. doi:10.1038/s41584-024-01130-8 PubMed PMID: 39014254.

15. Miao MZ, Su QP, Cui Y, Bahnson EM, Li G, Wang M, et al. Redox-active endosomes mediate α5β1 integrin signaling and promote chondrocyte matrix metalloproteinase production in osteoarthritis. Sci Signal. 2023;16(809). doi:10.1126/scisignal.adf8299 PubMed PMID: 37906629.

16. Gerwin N, Scotti C, Halleux C, Fornaro M, Elliott J, Zhang Y, et al. Angiopoietin-like 3-derivative LNA043 for cartilage regeneration in osteoarthritis: a randomized phase 1 trial. Nat Med. 2022 Dec 1;28(12):2633–45. doi:10.1038/s41591-022-02059-9 PubMed PMID: 36456835.

17. Wee WKJ, Low ZS, Ooi CK, Henategala BP, Lim ZGR, Yip YS, et al. Single-cell analysis of skin immune cells reveals an Angptl4-ifi20b axis that regulates monocyte differentiation during wound healing. Cell Death Dis. 2022 Feb 1;13(2). doi:10.1038/s41419-022-04638-7 PubMed PMID: 35210411.

18. Huang W, Jiang L, Jiang Y, Li S, Liu W, Zong K, et al. ANGPTL4 induces Kupffer cell M2 polarization to mitigate acute rejection in liver transplantation. Sci Rep. 2025 Dec 1;15(1). doi:10.1038/s41598-024-81832-x PubMed PMID: 39762255.

19. Schumacher A, Denecke B, Braunschweig T, Stahlschmidt J, Ziegler S, Brandenburg LO, et al. Angptl4 is upregulated under inflammatory conditions in the bone marrow of mice, expands myeloid progenitors, and accelerates reconstitution of platelets after myelosuppressive therapy. J Hematol Oncol. 2015 Dec 12;8(1). doi:10.1186/s13045-015-0152-2 PubMed PMID: 26054961.

20. Sylvers-Davie KL, Davies BSJ. Regulation of lipoprotein metabolism by ANGPTL3, ANGPTL4, and ANGPTL8. American Journal of Physiology - Endocrinology and Metabolism. American Physiological Society; 2021. p. E493–508. doi:10.1152/AJPENDO.00195.2021 PubMed PMID: 34338039.

21. Yang L, Wang Y, Xu Y, Li K, Yin R, Zhang L, et al. ANGPTL3 is a novel HDL component that regulates HDL function. J Transl Med. 2024 Dec 1;22(1). doi:10.1186/s12967-024-05032-x PubMed PMID: 38462608.

22. Chen PY, Gao WY, Liou JW, Lin CY, Wu MJ, Yen JH. Angiopoietin-like protein 3 (Angptl3) modulates lipoprotein metabolism and dyslipidemia. International Journal of Molecular Sciences. MDPI; 2021. doi:10.3390/ijms22147310 PubMed PMID: 34298929.

23. Kersten S. Angiopoietin-like 3 in lipoprotein metabolism. Nature Reviews Endocrinology. Nature Publishing Group; 2017. p. 731–9. doi:10.1038/nrendo.2017.119 PubMed PMID: 28984319.

24. Biterova E, Esmaeeli M, Alanen HI, Saaranen M, Ruddock LW. Structures of Angptl3 and Angptl4, modulators of triglyceride levels and coronary artery disease. Sci Rep. 2018 Dec 1;8(1). doi:10.1038/s41598-018-25237-7 PubMed PMID: 29713054.

25. Camenisch G, Pisabarro MT, Sherman D, Kowalski J, Nagel M, Hass P, et al. ANGPTL3 stimulates endothelial cell adhesion and migration via integrin αvβ3 and induces blood vessel formation in vivo. Journal of Biological Chemistry. 2002 May 10;277(19):17281–90. doi:10.1074/jbc.M109768200 PubMed PMID: 11877390.

26. Zhang Y, Yan C, Dong Y, Zhao J, Yang X, Deng Y, et al. ANGPTL3 accelerates atherosclerotic progression via direct regulation of M1 macrophage activation in plaque. J Adv Res. 2025 Apr 1;70:125–38. doi:10.1016/j.jare.2024.05.011 PubMed PMID: 38740260.

27. Ma Y, Chen Y, Xu H, Du N. The influence of angiopoietin-like protein 3 on macrophages polarization and its effect on the podocyte EMT in diabetic nephropathy. Front Immunol. 2023;14. doi:10.3389/fimmu.2023.1228399 PubMed PMID: 37638046.

28. Clark K, Pankov R, Travis MA, Askari JA, Mould AP, Craig SE, et al. A specific α5β1-integrin conformation promotes directional integrin translocation and fibronectin matrix formation. J Cell Sci. 2005 Jan 15;118(2):291–300. doi:10.1242/jcs.01623 PubMed PMID: 15615773.

29. Miao MZ, Su QP, Cui Y, Bahnson EM, Li G, Wang M, et al. Redox-active endosomes mediate α5β1 integrin signaling and promote chondrocyte matrix metalloproteinase production in osteoarthritis. Sci Signal. 2023;16(809). doi:10.1126/scisignal.adf8299 PubMed PMID: 37906629.

30. Maylin AB, Irene GLl, Villarroya O, María BJ, Kouri JB, Costell M. Genetic abrogation of the fibronectin-α5β1 integrin interaction in articular cartilage aggravates osteoarthritis in mice. PLoS One. 2018 Jun 1;13(6). doi:10.1371/journal.pone.0198559 PubMed PMID: 29870552.

31. Takano M, Hirose N, Sumi C, Yanoshita M, Nishiyama S, Onishi A, et al. ANGPTL2 Promotes Inflammation via Integrin α5β1 in Chondrocytes. Cartilage. 2021 Dec 1;13(2_suppl):885S–897S. doi:10.1177/1947603519878242 PubMed PMID: 31581797.

32. Martino MM, Hubbell JA. The 12th–14th type III repeats of fibronectin function as a highly promiscuous growth factor-binding domain. The FASEB Journal. 2010 Dec;24(12):4711–21. doi:10.1096/fj.09-151282 PubMed PMID: 20671107.

33. Wijelath ES, Rahman S, Namekata M, Murray J, Nishimura T, Mostafavi-Pour Z, et al. Heparin-II domain of fibronectin is a vascular endothelial growth factor-binding domain: Enhancement of VEGF biological activity by a singular growth factor/matrix protein synergism. Circ Res. 2006 Oct;99(8):853–60. doi:10.1161/01.RES.0000246849.17887.66 PubMed PMID: 17008606.

34. Schumacher S, Dedden D, Vazquez Nunez R, Matoba K, Takagi J, Biertümpfel C, et al. Structural insights into integrin α 5 β 1 opening by fibronectin ligand. Sci. Adv [Internet]. 2021. Available from: https://www.science.org

35. Bachmann M, Kukkurainen S, Hytönen VP, Wehrle-Haller B. CELL ADHESION BY INTEGRINS. Physiol Rev. 2019;99:1655–99. doi:10.1152/physrev.00036.2018.-Integrins

36. Chastney MR, Kaivola J, Leppänen VM, Ivaska J. The role and regulation of integrins in cell migration and invasion. Nature Reviews Molecular Cell Biology. Nature Research; 2025. p. 147–67. doi:10.1038/s41580-024-00777-1 PubMed PMID: 39349749.

37. Utomo L, Bastiaansen-Jenniskens YM, Verhaar JAN, van Osch GJVM. Cartilage inflammation and degeneration is enhanced by pro-inflammatory (M1) macrophages in vitro, but not inhibited directly by anti-inflammatory (M2) macrophages. Osteoarthritis Cartilage. 2016 Dec 1;24(12):2162–70. doi:10.1016/j.joca.2016.07.018 PubMed PMID: 27502245.

38. Garcia J, Hulme C, Mennan C, Roberts S, Bastiaansen-Jenniskens YM, van Osch GJVM, et al. The synovial fluid from patients with focal cartilage defects contains mesenchymal stem/stromal cells and macrophages with pro- and anti-inflammatory phenotypes. Osteoarthr Cartil Open. 2020 Jun 1;2(2). doi:10.1016/j.ocarto.2020.100039

39. Panichi V, Costantini S, Grasso M, Arciola CR, Dolzani P. Innate Immunity and Synovitis: Key Players in Osteoarthritis Progression. International Journal of Molecular Sciences. Multidisciplinary Digital Publishing Institute (MDPI); 2024. doi:10.3390/ijms252212082 PubMed PMID: 39596150.

40. Ogle ME, Segar CE, Sridhar S, Botchwey EA. Monocytes and macrophages in tissue repair: Implications for immunoregenerative biomaterial design. Experimental Biology and Medicine. SAGE Publications Inc.; 2016. p. 1084–97. doi:10.1177/1535370216650293 PubMed PMID: 27229903.

41. Shan W, Cheng C, Huang W, Ding Z, Luo S, Cui G, et al. Angiopoietin-like 2 upregulation promotes human chondrocyte injury via NF-κB and p38/MAPK signaling pathway. J Bone Miner Metab. 2019 Nov 1;37(6):976–86. doi:10.1007/s00774-019-01016-w PubMed PMID: 31214838.

42. Schumacher A, Denecke B, Braunschweig T, Stahlschmidt J, Ziegler S, Brandenburg LO, et al. Angptl4 is upregulated under inflammatory conditions in the bone marrow of mice, expands myeloid progenitors, and accelerates reconstitution of platelets after myelosuppressive therapy. J Hematol Oncol. 2015 Dec 12;8(1). doi:10.1186/s13045-015-0152-2 PubMed PMID: 26054961.

43. Li X, Zhang Y, Zhang M, Wang Y. GALNT2 regulates ANGPTL3 cleavage in cells and in vivo of mice. Sci Rep. 2020 Dec 1;10(1). doi:10.1038/s41598-020-73388-3 PubMed PMID: 32999434.

44. Schjoldager KTBG, Vester-Christensen MB, Bennett EP, Levery SB, Schwientek T, Yin W, et al. O-glycosylation modulates proprotein convertase activation of angiopoietin-like protein 3: Possible role of polypeptide GalNAc-transferase-2 in regulation of concentrations of plasma lipids. Journal of Biological Chemistry. 2010 Nov 19;285(47):36293–303. doi:10.1074/jbc.M110.156950 PubMed PMID: 20837471.

45. Hao C, Zou Q, Bai X, Shi W. Effect of glycosylation on protein folding: From biological roles to chemical protein synthesis. iScience. Elsevier Inc.; 2025. doi:10.1016/j.isci.2025.112605

46. He M, Zhou X, Wang X. Glycosylation: mechanisms, biological functions and clinical implications. Signal Transduction and Targeted Therapy. Springer Nature; 2024. doi:10.1038/s41392-024-01886-1 PubMed PMID: 39098853.

47. Tanoue H, Morinaga J, Yoshizawa T, Yugami M, Itoh H, Nakamura T, et al. Angiopoietin-like protein 2 promotes chondrogenic differentiation during bone growth as a cartilage matrix factor. Osteoarthritis Cartilage. 2018 Jan 1;26(1):108–17. doi:10.1016/j.joca.2017.10.011 PubMed PMID: 29074299.

48. Murata M, Yudo K, Nakamura H, Chiba J, Okamoto K, Suematsu N, et al. Hypoxia upregulates the expression of angiopoietin-like-4 in human articular chondrocytes: Role of angiopoietin-like-4 in the expression of matrix metalloproteinases and cartilage degradation. Journal of Orthopaedic Research. 2009 Jan;27(1):50–7. doi:10.1002/jor.20703 PubMed PMID: 18634015.

49. Jia C, Li X, Pan J, Ma H, Wu D, Lu H, et al. Silencing of Angiopoietin-Like Protein 4 (Angptl4) Decreases Inflammation, Extracellular Matrix Degradation, and Apoptosis in Osteoarthritis via the Sirtuin 1/NF-κ B Pathway. Oxid Med Cell Longev. 2022;2022. doi:10.1155/2022/1135827 PubMed PMID: 36071864.

50. Wang JY, Liu YH, Wang X, Ma M, Pan ZY, Fan AY, et al. Atf3 + senescent chondrocytes mediate meniscus degeneration in aging. Arthritis Res Ther. 2025 Dec 1;27(1). doi:10.1186/s13075-025-03566-z PubMed PMID: 40375324.

51. Jumper J, Evans R, Pritzel A, Green T, Figurnov M, Ronneberger O, et al. Highly accurate protein structure prediction with AlphaFold. Nature. 2021 Aug 26;596(7873):583–9. doi:10.1038/s41586-021-03819-2 PubMed PMID: 34265844.

52. Li H, Qiu J, Fu XD. RASL-seq for Massively Parallel and Quantitative Analysis of Gene Expression. Curr Protoc Mol Biol. 2012 Apr;1(SUPPL.98). doi:10.1002/0471142727.mb0413s98 PubMed PMID: 22470064.

53. Ritchie ME, Phipson B, Wu D, Hu Y, Law CW, Shi W, et al. Limma powers differential expression analyses for RNA-sequencing and microarray studies. Nucleic Acids Res. 2015 Jan 6;43(7):e47. doi:10.1093/nar/gkv007 PubMed PMID: 25605792.

54. Korotkevich G, Sukhov V, Budin N, Shpak B, Artyomov MN, Sergushichev A. Fast gene set enrichment analysis [Internet]. 2016. Available from: http://biorxiv.org/lookup/doi/10.1101/060012 doi:10.1101/060012

