## Supplementary Materials for "LNA043 and ANGPTL3 Interactions with Integrin α5β1 and Fibronectin Drive Distinct Stromal and Immune Responses"

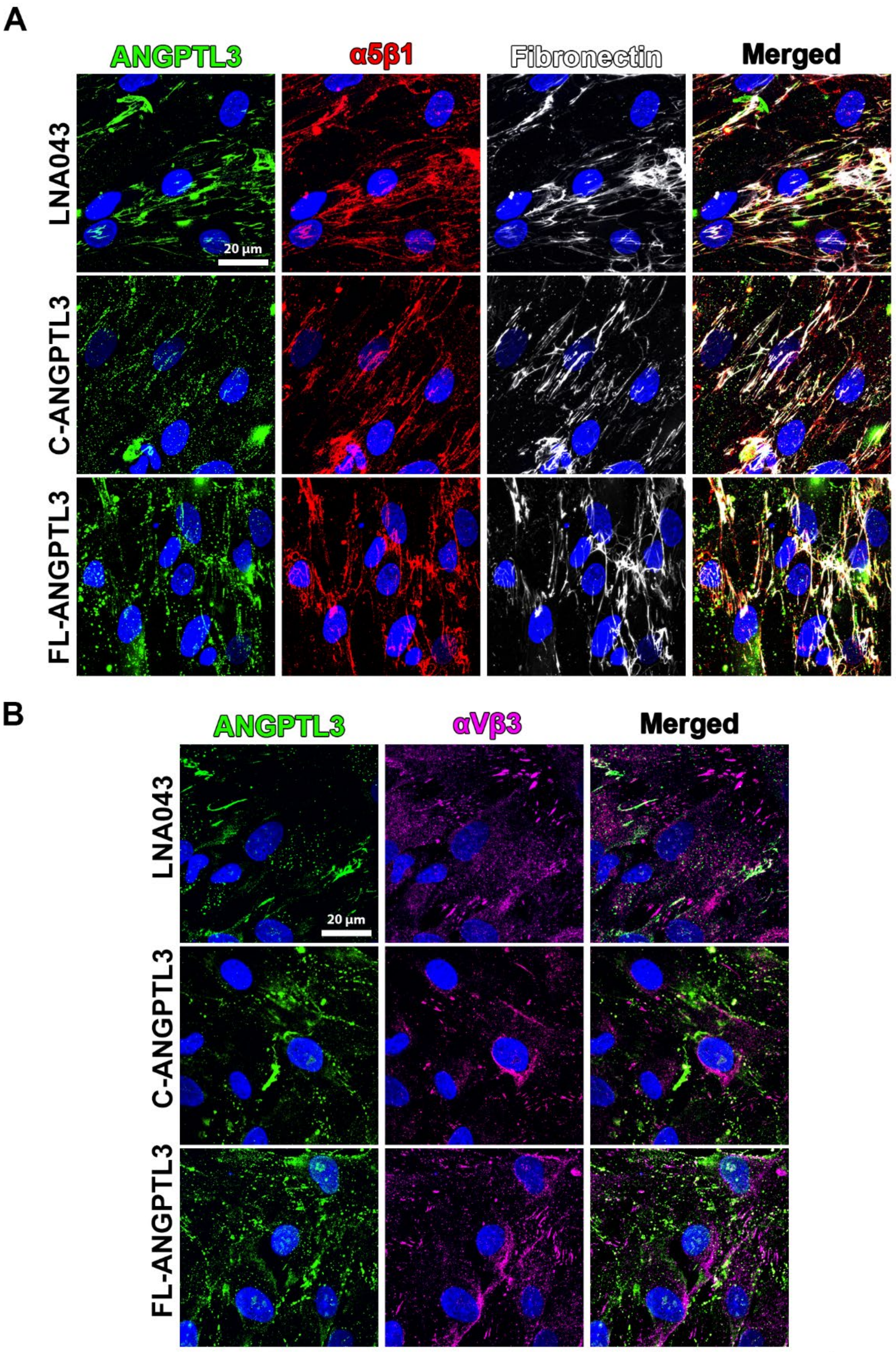


**Figure Supplementary 1**

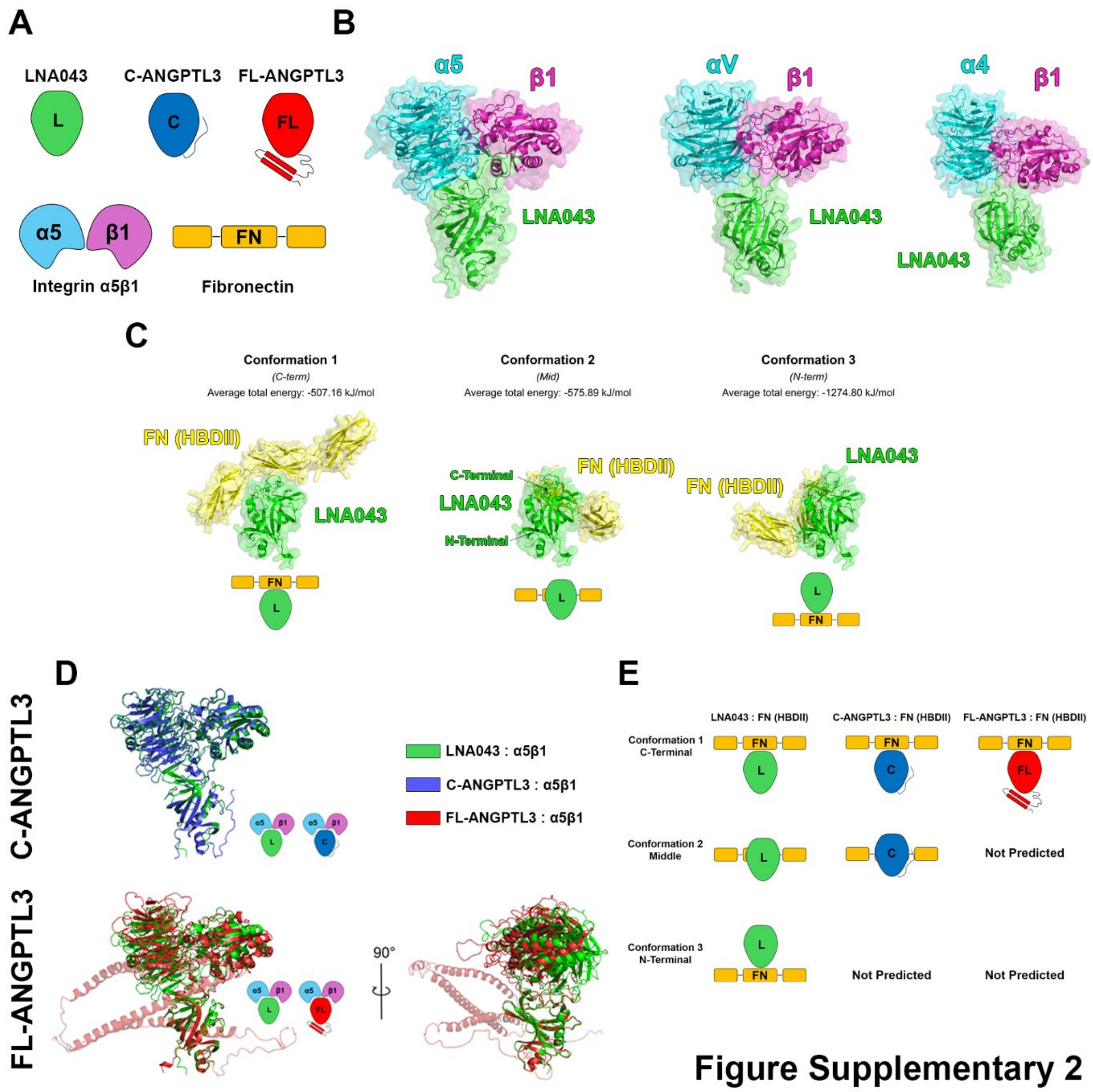


**Figure Supplementary 2**

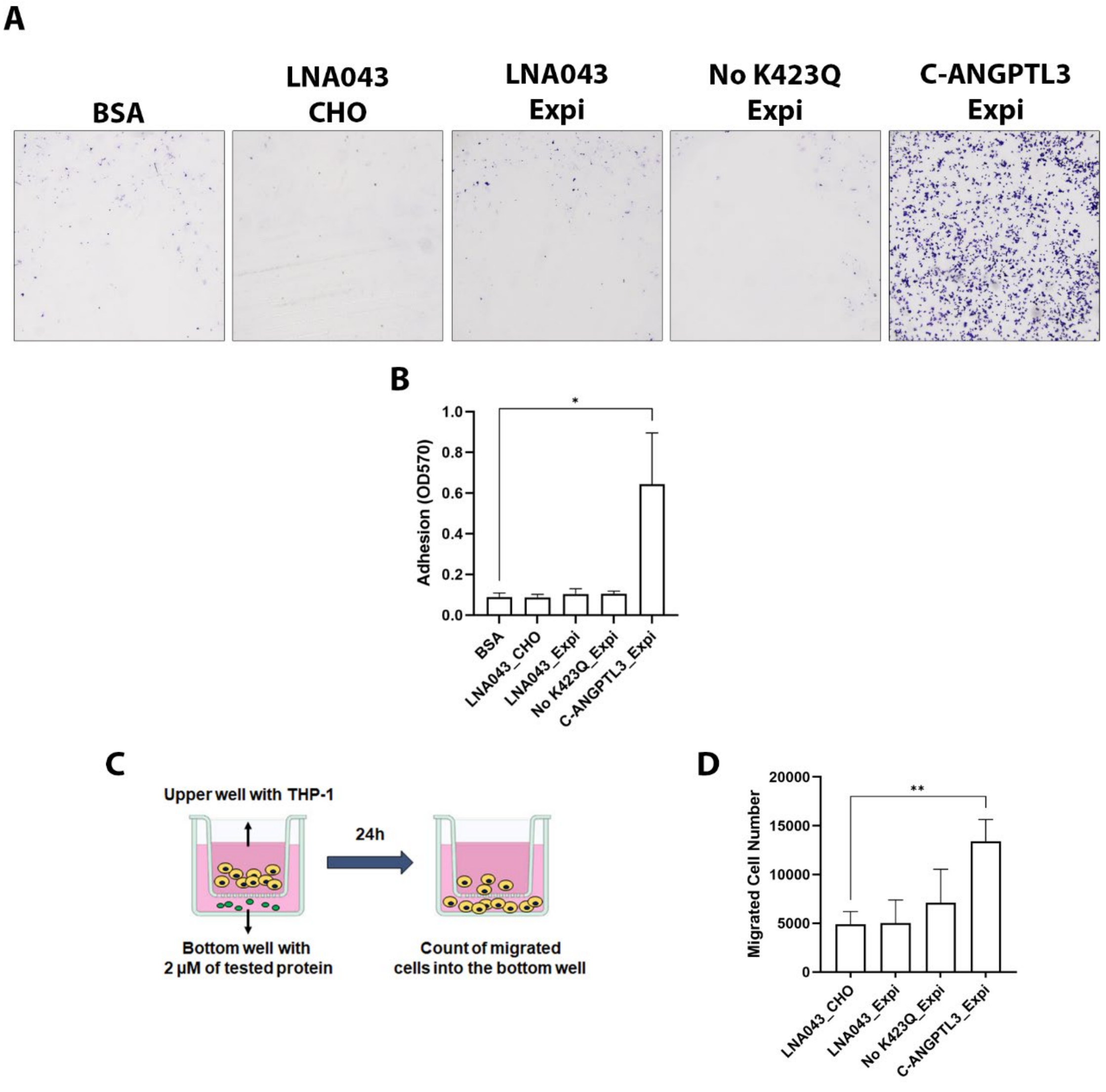


#### Figure Supplementary 3

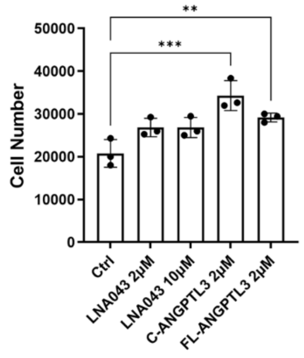


#### Figure Supplementary 4

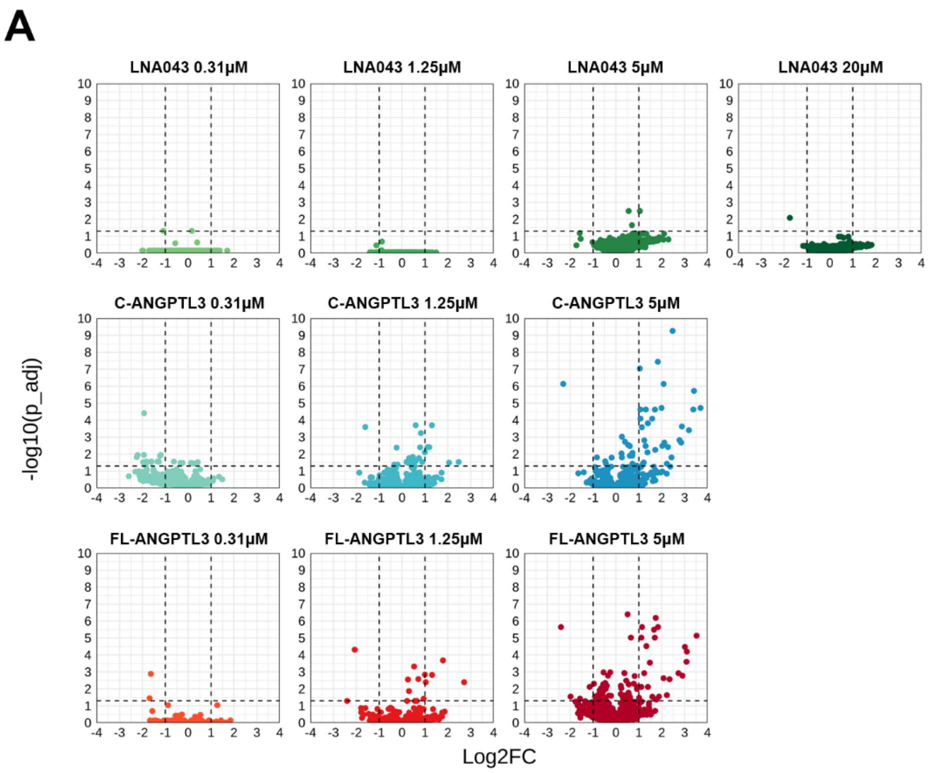


### Figure Supplementary 5

| Ligand | Binder | Binding confor mation | Average Coulombic energy (kJ/mol) | Average Lenard-Jones energy (kJ/mol) | Average total energy (kJ/mol) | RMSD total energy (kJ/mol) |
|---|---|---|---|---|---|---|
| LNA043 | α5β1 | - | -1505.46 | -470.04 | -1975.50 | 246.79 |
| LNA043 | αVβ1 | - | -2038.45 | -138.06 | -2176.51 | 230.82 |
| LNA043 | α4β1 | - | -892.88 | -290.53 | -1183.41 | 167.90 |
| LNA043 | FN 12-14 (HBDII) | 1 | -233.42 | -273.74 | -507.16 | 121.58 |
| LNA043 | FN 12-14 (HBDII) | 2 | -442.52 | -133.37 | -575.89 | 236.63 |
| LNA043 | FN 12-14 (HBDII) | 3 | -1082.28 | -192.51 | -1274.80 | 293.54 |
| C-ANGPTL3 | α5β1 | - | -1398.70 | -376.57 | -1775.27 | 157.49 |
| C-ANGPTL3 | FN 12-14 (HBDII) | 1 | -297.81 | -198.48 | -496.29 | 271.00 |
| C-ANGPTL3 | FN 12-14 (HBDII) | 2 | -582.00 | -163.76 | -745.76 | 142.98 |
| FL-ANGPTL3 | α5β1 | - | -1597.23 | -261.78 | -1859.01 | 238.88 |
| FL-ANGPTL3 | FN 12-14 (HBDII) | 1 | -963.71 | -274.34 | -1238.042 | 314.13 |

### Supplementary Table

**Figure Supplementary 2. Multi-Partner Binding of LNA043 and ANGPLT3: Structural Predictions and Cellular Validation.** (A) Schematic representation of structures used in the study. (B) Predicted AlphaFold structures of LNA043 in complex with integrin α5β1 (left), αVβ1 (center) and α4β1 (right). (C) Predicted conformations of LNA043 : FN HBDII complex. The calculated total average energy is shown at the top of each conformation. (D) Comparison of predicted α5β1 complex. LNA043 : α5β1 (green) is aligned with C-ANGPTL3 : α5β1 (blue) and with FL-ANGPTL3 : α5β1 (red). The coil-coiled domain of FL-ANGPTL3 is displayed as semitransparent since it does not participate in the binding and it has a random relative orientation. (E) Comparison of predicted complex with fibronectin HBDII. Conformations are named based on which area of the protein is involved in the binding: C-terminal area (top), middle (center) or N-terminal area (bottom). When a specific conformation was not predicted by AlphaFold, it is listed as not predicted.

**Figure Supplementary 3. Assessment of the impact of K423Q point mutation or cell production system on LNA043 potency.** (A) Representative images of ihMSCs adherent to wells coated with 2 μM of LNA043 produced in CHO, LNA043 produced in Expi293 with (K243Q) or without point mutation (No K423Q), and C-ANGPTL3 after 1 hour at 37 °C. BSA coated wells were used as negative control. (B) Quantification of ihMSC adhesion. (C) Schematic of transwell migration assay. THP-1 cells were seeded in the upper chamber and 2 μM of tested proteins were added to the lower chamber. Migration was assessed after 24 hours. (D) Quantification of the number of migrated cells in the bottom well after 24 hours. Data are shown as mean ± SD from 3 independent experiments. *$p < 0.05$. **$p < 0.01$.

**Figure Supplementary 4. THP-1 proliferation in response to LNA043, C-ANGPTL3 or FL-ANGPTL3.** Measurement of THP-1 cell numbers following 24 hours incubation with 2 or 10 μM LNA043, 2 μM C-ANGPTL3, or FL-ANGPTL3. Data are shown as mean ± SD from 3 independent experiments. **$p<0.01$, ***$p<0.001$.

**Figure Supplementary 5. Transcriptomic analysis of THP-1 treated with increasing doses of LNA043, C-ANGPTL3 and FL-ANGPTL3.** (A) Volcano plots illustrating differentially expressed genes (DEGs) after 24 hours treatment with 0.31, 1.25, 5 and 20 μM of LNA043 (green), 0.31, 1.25, 5 μM C-ANGPTL3 (cyan) or FL-ANGPTL3 (Orange). Data are expressed as log2 fold change (LOG2FC, x-axis) relative to untreated control, and significance is expressed as -log10 adjusted p-value (-log10(p_adj), y axis). Genes are considered significantly regulated above thresholds of LOG2FC ≥ 1 or ≤ –1 and –log10(p_adj) ≥ 1.3 (p_adj < 0.05).

**Supplementary Table** . List of average interacting energies (Coulombic, Lenard-Jones or total) between ANGPTL3 variants and integrins or fibronectin HBDII domain.

**Supplementary Video .** Mapping LNA043 molecular interactions with integrin α5β1 and Fibronectin (FN). The model of the predicted ternary complex is displayed with structures of LNA043 (green), integrin α5β1 (cyan/magenta) and Fibronectin HBDII domain (yellow) in a cartoon and semitransparent surface representation. A schematic representation is depicted on top to facilitate the visualization. Motions are artificially set up to illustrate binding, and do not represent realistic molecular dynamics.
